# Electrical match between initial segment and somatodendritic compartment for action potential backpropagation in retinal ganglion cells

**DOI:** 10.1101/2020.09.15.297937

**Authors:** Sarah Goethals, Martijn C. Sierksma, Xavier Nicol, Annabelle Réaux-Le Goazigo, Romain Brette

## Abstract

The action potential of most vertebrate neurons initiates in the axon initial segment (AIS), and is then transmitted to the soma where it is regenerated by somatodendritic sodium channels. For successful transmission, the AIS must produce a strong axial current, so as to depolarize the soma to the threshold for somatic regeneration. Theoretically, this axial current depends on AIS geometry and Na^+^ conductance density. We measured the axial current of mouse RGCs using whole-cell recordings with post-hoc AIS labeling. We found that this current is large, implying high Na^+^ conductance density, and carries a charge that co-varies with capacitance so as to depolarize the soma by ~30 mV. Additionally, we observed that the axial current attenuates strongly with depolarization, consistent with sodium channel inactivation, but temporally broadens so as to preserve the transmitted charge. Thus, the AIS appears to be organized so as to reliably backpropagate the axonal action potential.

## Introduction

In most vertebrate neurons, action potentials (APs) initiate in the axon initial segment (AIS), a highly organized structure near the soma (Bender & Trussell, 2012), then propagate forward to the axon terminals and backward to the soma and dendrites (Debanne *et al.*, 2011). This backward transmission is functionally important for synaptic plasticity, which requires a precisely timed signal of firing activity at the synapse (Caporale & Dan, 2008). It is also important for long-term intrinsic plasticity since the soma holds the genetic material (Daoudal & Debanne, 2003), and presumably also for structural plasticity of the AIS, which depends on somatic voltage-gated calcium channels (Evans *et al.*, 2013).

At spike initiation, the soma receives an axial current from the AIS, which depolarizes the membrane. When the somatic membrane potential is depolarized about 30 mV above firing threshold, the AP is regenerated by somatic sodium channels (Kole & Stuart, 2008). Hamada et al. (2016) found indirect evidence that the axial current matches the capacitance of the somatodendritic compartment, as they observed that larger cortical pyramidal cells tend to have a more proximal AIS, which should theoretically produce a stronger current. However, the axial current was not directly measured.

Such a measurement could also allow estimating the conductance density of AIS sodium channels, in particular using resistive coupling theory (Brette, 2013; Kole & Brette, 2018). The fact that the AIS, a small structure, must produce a current able to charge a much larger piece of membrane (soma and proximal dendrites), suggests that conductance density is high, in agreement with immunochemical observations (Lorincz & Nusser, 2010). However, this has remained a somewhat contentious issue (Fleidervish *et al.*, 2010) because direct patch-clamp measurements in the intact AIS indicate low conductance density (Colbert & Pan, 2002), which could be an artifact due to the anchoring of channels to the cytoskeleton (Kole *et al.*, 2008).

Finally, it is known that sodium channels can inactivate substantially below threshold, resulting in spike threshold adaptation (Azouz & Gray, 2000; Platkiewicz & Brette, 2011; Fontaine *et al.*, 2014). This suggests that the axial current at spike initiation may also vary substantially. If this is the case, then how can spikes be reliably transmitted to the soma?

To address these questions, we measured the axial current and spontaneous action potentials in ganglion cells of isolated mouse retina followed by ankyrin-G-antibody labeling to measure AIS geometry. We examined the axial current at spike initiation, just below threshold, and with threshold adaptation, and compared these results to theoretical predictions.

## Results

### The axial current at spike initiation

#### Action potentials of retinal ganglion cells

Early work on vertebrate motoneurons showed that APs recorded in the soma typically consist of two components (Fatt, 1957; Coombs *et al.*, 1957): an abrupt depolarization due to the axial current originating from the axon initial segment, followed by a regeneration of the AP at a higher potential, by the opening of somatic sodium channels. These two components are clearly distinguished in recordings of spontaneous APs of retinal ganglion cells (RGC). Figure 1A illustrates a typical AP, rising abruptly from threshold. The voltage derivative *dV/dt* shows two distinct components, which appear most clearly when plotted against the membrane potential *V* (Fig. 1B). We define the regeneration threshold as the potential when the acceleration *d^2^V/dt^2^* is maximal. In n = 10 cells (see Methods for the inclusion criteria), we observed that the spike threshold of spontaneous APs was −49 ± 3.8 mV (s.d.) while the regeneration threshold is −16 ± 4.6 mV (s.d.), about 33 ± 5 mV (s.d.) higher (Fig. 1C). This is similar to previous measurements in layer 5 cortical pyramidal cells (Kole & Stuart, 2008).

**Figure 1.**
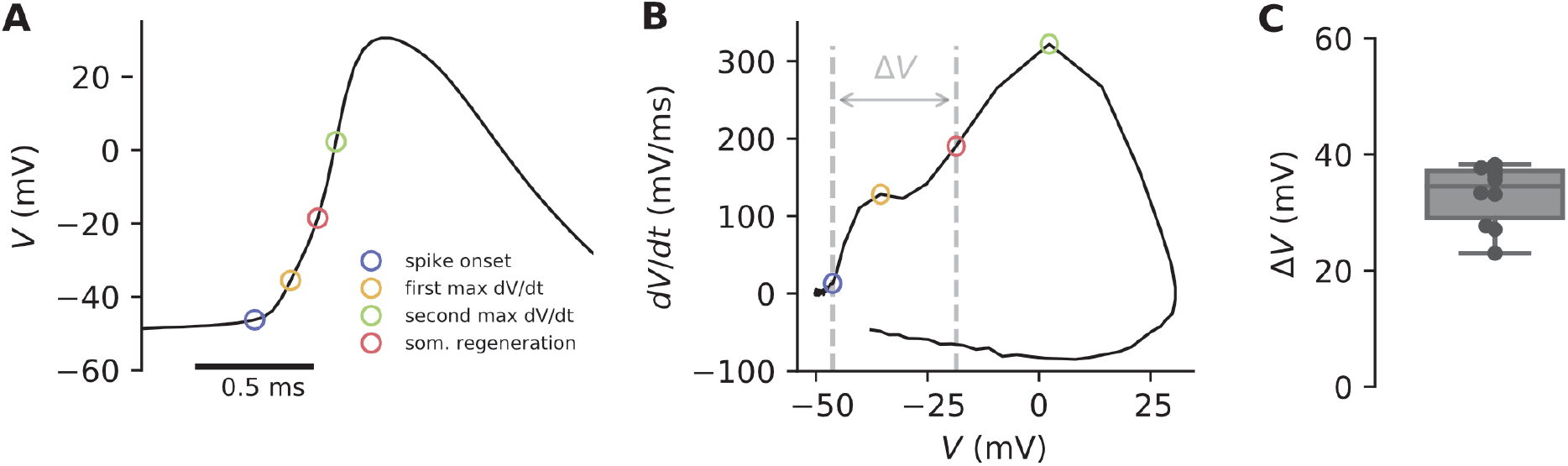
Spontaneous APs of retinal ganglion cells consist of two components. A, Spontaneous AP of a RGC, highlighting spike onset (blue), point of maximum dV/dt of the first component (yellow), somatic regeneration threshold (red), point of maximum dV/dt of the second component (green). B, Phase plot showing dV/dt vs. V. The double arrow shows the difference ∆V between spike onset and regeneration threshold. C, Statistics of ∆V.

#### Transmission of the axial current to the soma

We measured the axial current with whole-cell voltage clamp by stepping the command potential from V0 = −60 mV to a variable potential *V*. Voltage steps above a threshold value evoke large spikes of inward current (Fig. 2A). When the peak current is plotted against voltage, a sharp discontinuity is seen (Fig. 2B). Similar recordings have been reported in several cell types in whole-cell patch (Magistretti *et al.*, 2006; Diwakar *et al.*, 2009; Milescu *et al.*, 2010), and also in two-electrode voltage clamp recordings of cat motoneurons (Barrett & Crill, 1980). As argued by Milescu et al. (2010), this abrupt increase in current most likely reflects the axial current produced by the AIS AP. Indeed, the current-voltage curve shows a plateau reflecting the all-or-none axonal spike, followed by an increase at higher potential, most likely reflecting the somatic sodium current. These currents were eliminated by 1 μM tetrodotoxin, a potent sodium channel blocker (2 ±0.4 % current remaining, *n* = 4 cells, paired t_3_ = 4.5, p = 0.02). For further analysis, the current measurements were corrected for the errors introduced by the series resistance, as described by Traynelis (1998) (see Methods).

**Figure 2.**
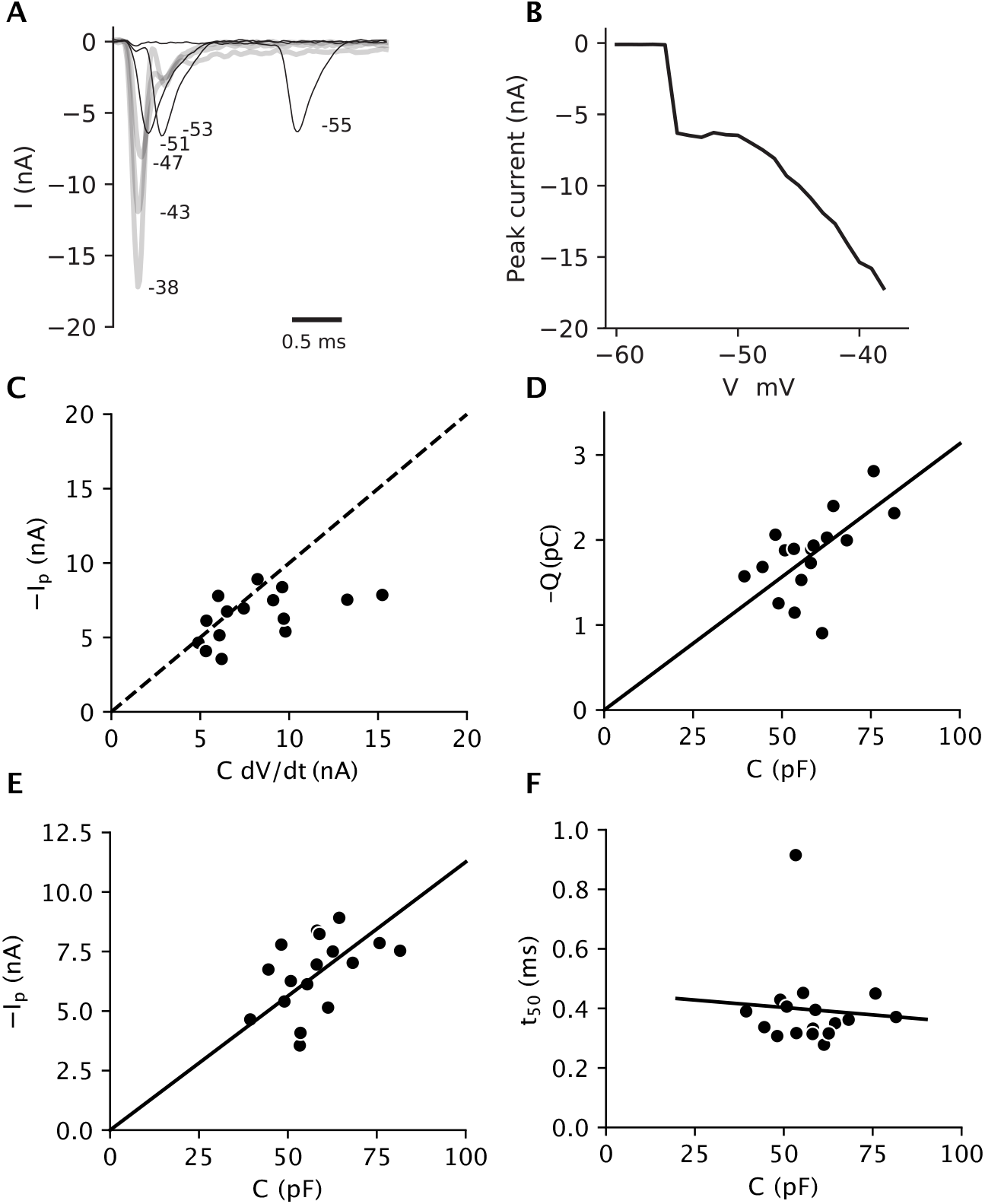
Transmission of the axial current to the soma. A, Raw current recordings (with leak current correction, and only online correction series resistance) in response to voltage steps from −60 mV to different target potentials (−55 mV to −38 mV). B, Peak current vs. step command potential. The peak current increases abruptly when the potential exceeds some threshold. C, Corrected peak axial current at spike initiation measured in voltage clamp vs. capacitive current measured in current clamp (n = 15; 5 cells were excluded because spontaneous APs or capacitance were not measured). The dashed diagonal line represents the identity −I_p_ = CdV/dt (not a regression line). D, Total transmitted charge vs. somatic capacitance (n = 17). The solid line is the best linear fit, with slope 31 mV. C, Peak axial current measured in voltage clamp vs. somatic capacitance measured in current clamp (n = 17). The solid line is the best linear fit (not affine; i.e., of the form −I_p_ = a.C). E, Axial current duration t_50_ measured at 50% of peak current, vs. capacitance, with regression line (n= 17); t_50_ = 0.36 ± 0.05 ms, excluding one clear outlier.

When the soma is not voltage-clamped, the axial current at spike initiation charges the somatic capacitance (and proximal dendrites). Therefore, we expect that the axial current measured in voltage clamp is approximately equal to the capacitive current *C.dV/dt* during the initial rise of an AP recorded in current clamp. We estimated the effective capacitance on the first ms of the response to a small current pulse, and measured *dV/dt* in the initial phase of a spontaneous AP (see Methods). We found that for most cells, the axial current measured in voltage clamp was indeed close to the capacitive current of a spontaneous AP (Fig. 2C). The somatic depolarization due to the axial current is Δ*V* = *Q*/*C*, where *Q* is the total charge *Q* transmitted to the soma, i.e., the integral of the axial current. We found a linear correlation between *Q* and *C* (Fig. 2D; Pearson correlation r = 0.56, p = 0.02), with a slope Δ*V* = 31 mV. This is remarkably close to the difference between spike threshold and regeneration threshold we observed on spontaneous APs (33 mV; Fig. 1C). Thus, the transmitted charge appears to be just enough to bring the somatic potential to the voltage where somatic sodium channels open and regenerate the AP.

This correlation was likely due to a linear relation between axial current and capacitance, because axial current increased with capacitance (Fig. 2E) while current duration was essentially constant (Fig. 2F). However, the relation with current was statistically less strong than with charge (Pearson correlation r= 0.47, p = 0.06), possibly because the measurement of *Q* is more reliable than that of *I_p_*, as integration reduces noise and charge is not affected by electrode filtering.

#### Axial current and AIS geometry

Theoretically, axial current depends on AIS geometry (Hamada *et al.*, 2016). We measured the geometry of the AIS of n = 14 cells by immunolabeling ankyrin-G, while identifying recorded cells using biocytin in the patch pipette (see Methods) (Fig. 3A). The AIS was on average 31 μm long (± 6 μm s.d.) and started at 8.6 ± 3.3 μm from the soma (Fig. 3B), with no statistically significant correlation between the two measurements (p = 0.59, Pearson test). We also measured the axon diameter at the proximal and distal ends of the AIS. Keeping in mind that these measurements approach the limits of conventional light microscopy, the proximal and distal diameters were 0.9 ± 0.3 μm and 0.5 ± 0.2 μm, respectively. For comparison, Raghuram et al. (2019) found by a similar method 1 μm and 0.6 μm on average in α S RGCs of mice, with substantial variability.

**Figure 3.**
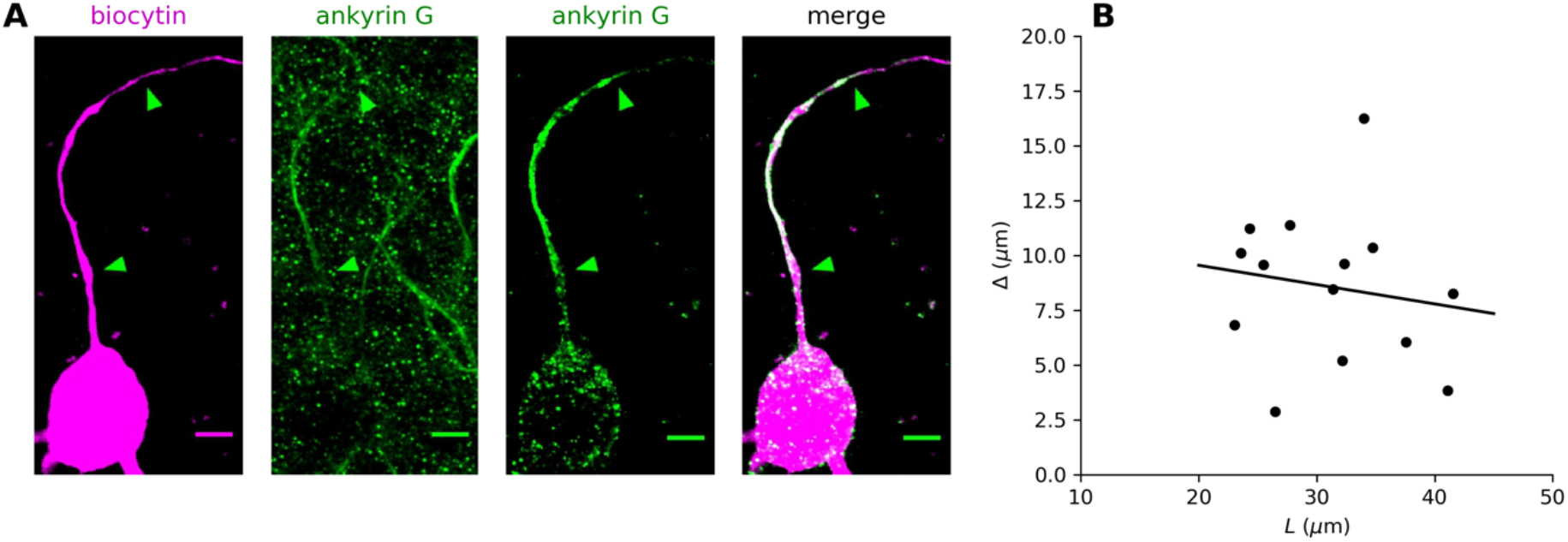
Geometry of the AIS. A left, Fluorescence image of a RGC labelled with biocytin (pink). The start and end position of the AIS are indicated by green arrowheads. Note that the biocytin staining extends beyond the AIS. Scale bar is 5 μm. A middle left, Fluorescence image of AISs labeled with ankyrin-G antibodies (green). A middle right, Fluorescence image of the AIS masked by the neuron morphology. A right, Merge of the first and third panels. B, Distance Δ of the AIS from the soma vs. AIS length L, with the regression line (p = 0.59, Pearson test).

The axial current was on average *I_p_* = −6.7 nA (s.d. 1.8 nA). A conservative estimate of Na+ conductance density *g_min_* in the AIS can be estimated by noting that the axial current cannot be greater than the maximum current that all Na^+^ channels can pass. This maximum Na+ current is *G*(*E_Na_* − *V*), where E_Na_ ≈ 70 mV is the reversal potential of Na^+^, *G* is the total Na^+^ conductance and *V* ≈ −15 mV is the local membrane potential at which the current through a Na^+^ channel is maximal (based on Na^+^ channel properties measured at the AIS of cortical pyramidal cells (Kole *et al.*, 2008)). Therefore, a lower bound for the total Na^+^ conductance is *G*_*min*_ = −*I_p_*/(*E_Na_* − *V*) ≈ 79 nS. Using the high estimate of 1 μm for the diameter, the average AIS area was therefore 97 μm^2^ (± 19 μm^2^ s.d.), which implies a minimum conductance density of 814 S/m^2^. With a diameter of 0.7 μm, we find a minimum of 1159 S/m^2^.

This is a conservative estimate because it ignores the conditions of propagation of the action potential. Resistive coupling theory provides a quantitative estimate of the axial current as a function of AIS geometry and *g*, by assuming that current entering the axonal membrane flows resistively towards the soma, which acts as a current sink (Brette, 2013; Hamada *et al.*, 2016; Kole & Brette, 2018; Goethals & Brette, 2020). Suppose first that conductance density is very high, such that the AIS is clamped at ENa when sodium channels open (Fig. 4A, dark green). Then by Ohm’s law, the AIS will produce an axial current *I_p_* = (*E_Na_*-*V_t_*)/*R_a_*, where *R_a_* is axial resistance between the soma and the proximal end of the AIS. Thus, we obtain an inverse relation between axial current and AIS position. However, conductance density is finite, which implies that the proximal side of the AIS is pulled towards the somatic potential (Fig. 4A, light green). This is electrically equivalent to shifting the AIS distally by an amount δ:

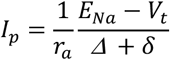

where Δ is the distance of the AIS from the soma, *r_a_* is the axial resistance per unit length, *V_t_* is the somatic spike threshold (*E_Na_* − *V*_*t*_ ≈ 120 mV) and

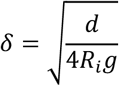

where *R_i_* is intracellular resistivity. Here, AIS length *L* can be neglected provided that it is substantially larger than δ (see Methods). This is clearly the case because *L* was 31 μm on average, while an upper estimate of δ using *d* = 1.2 μm and *g* = 1000 S/m^*2*^ is 17 μm. Thus, in our cells, AIS length should have no impact on axial current. The formula above agrees well with simulations of a simplified model with non-inactivating Na^+^ channels (Fig. 4B), except when the AIS is very proximal, where it gives an overestimation.

**Figure 4.**
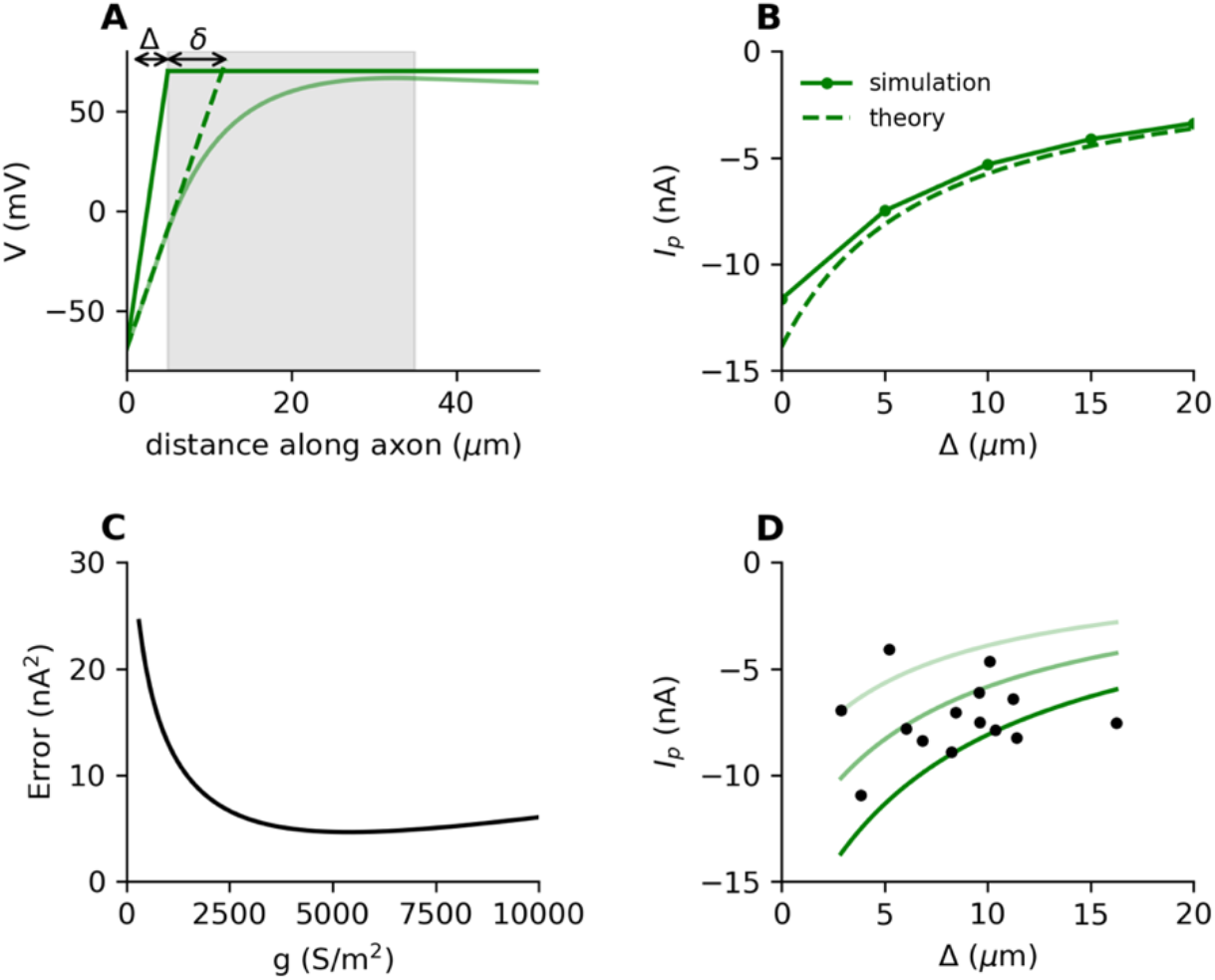
Predictions of axial current with resistive coupling theory. A, Light green curve: membrane potential along the axon of a simple model when all Na^+^ channels are open along the AIS, starting at distance Δ (gray shading), with a voltage clamp of the soma. Solid dark green line: idealized voltage profile for an AIS clamped at ENa. The axial current produced by the model is the same as the current produced by an AIS clamped at ENa and starting at distance Δ + δ (dashed line). B, Axial current at spike initiation vs. AIS position Δ in a simple model (solid) compared to theory (dashed). C, Mean squared error between predicted and measured current, as a function of conductance density g (with d = 1 μm). D, Measured axial current vs. Δ, with the theoretical relations for d = 0.8 μm, d = 1 μm and d = 1.2 μm (using g = 5400 S/m^2^, the minimum in C; n = 14).

This analysis shows that it is the proximal geometry of the AIS that matters for the calculation of the axial current. Using *d* = 1 μm, we find that the error between the predicted and the measured current varies with *g*, with a broad minimum at about 5500 S/m^2^ (Fig. 4C). This is close to the value that Guo et al. (2013) obtained by model optimization on current-clamp recordings (5000 S/m^2^). Figure 4D shows the axial current measured in our cells as a function of AIS position, together with the theoretical predictions using *g* = 5500 S/m^2^ with diameters *d* = 0.8 μm, *d* = 1 μm and *d* = 1.2 μm. Although the magnitude of the recorded axial currents is consistent with the theoretical prediction, the correlation between axial current and AIS position did not reach significance (*p* = 0.66, Pearson test). This may simply reflect the variability of AIS diameter, which has a strong impact on this relation.

This estimate relies on the precision of AIS geometry measurements. However, even if AIS position and length were different, high conductance density would still be necessary to account for the measured axial current. This can be shown by calculating the maximum axial current that can be generated across all possible AIS geometries for a given conductance density, and then deducing the minimum conductance density necessary to produce a given axial current *I* (see Methods). Taking *R_i_* = 100 Ω.cm, we find *g_min_* = 1263 S/m^2^ for *d* = 1 μm and *g_min_* = 2467 S/m^2^ for *d* = 0.8 μm (as argued above, it is mostly the geometry of the proximal side that matters for this calculation). With a higher value for *R*_*i*_, the lower bound on conductance density would be proportionally higher.

Overall, this analysis shows that the strong axial current produced at spike initiation requires a Na^+^ conductance density in the AIS of at least about 1000 S/m^2^ using the most conservative estimates, and plausibly several thousand S/m^2^ based on our measurements of AIS location and standard values of *R_i_*.

### The threshold axial current

#### Variation of axial current near threshold

In a model where the AIS is reduced to a single point, theory predicts that spikes initiate when the sodium current, and therefore the axial current, reaches a threshold *I*_*t*_ = *k*/*R_a_*, where *k* is the activation slope factor of sodium channels (*k* ≈ 5 mV) (Brette, 2013). This makes spike initiation distal from the soma efficient because the Na^+^ flux below threshold is low. We show in the Methods that the formula is approximately correct in an extended AIS model, if *R_a_* is measured between soma and the middle of the AIS. Thus, the threshold axial current is determined by AIS geometry. We tried to estimate *I_t_* in our cells.

To give an order of magnitude, with d = 1 μm and given that the middle position of the AIS is 24 μm on average, we obtain *R_a_* ≈ 31 MΩ, which gives *I_t_* ≈ 160 pA (assuming *k* = 5 mV), a small current. Figure 5A shows a recording of the axial current at threshold, which is noisy. We measure the peak current after smoothing. In addition, theory predicts that the axial current increases very steeply near threshold (*dI/dV* is infinite at threshold, see Methods), as shown in Figure 5B. This makes the threshold current difficult to measure, and likely leads to an underestimation of the threshold current. We measured the current at different step voltages in steps of 0.5 mV (*n* = 12). In the example shown in Fig. 5C, a small but noticeable current appears at 3 mV below threshold, which increases at higher subthreshold potentials. More precisely, theory predicts that *V-V_t_* is proportional to (I/I_t_ − 1)^2^. This relation is shown in a biophysical model in Figure 5D. In a simplified model (no sodium channel inactivation or potassium channels), the slope *β* is predicted to be equal to *k/2* (see Methods). This slope is found to be larger in the more realistic model shown in Figure 5D, *β* ≈ 4.7 mV, close to *k*. Our data fitted this quadratic relation well (Fig. 5E), with slopes *β* ≈ 4.2 mV (± 1.7 mV), in the expected range (Fig. 5F). Thus, the axial current increases steeply just below threshold, in agreement with theory.

**Figure 5.**
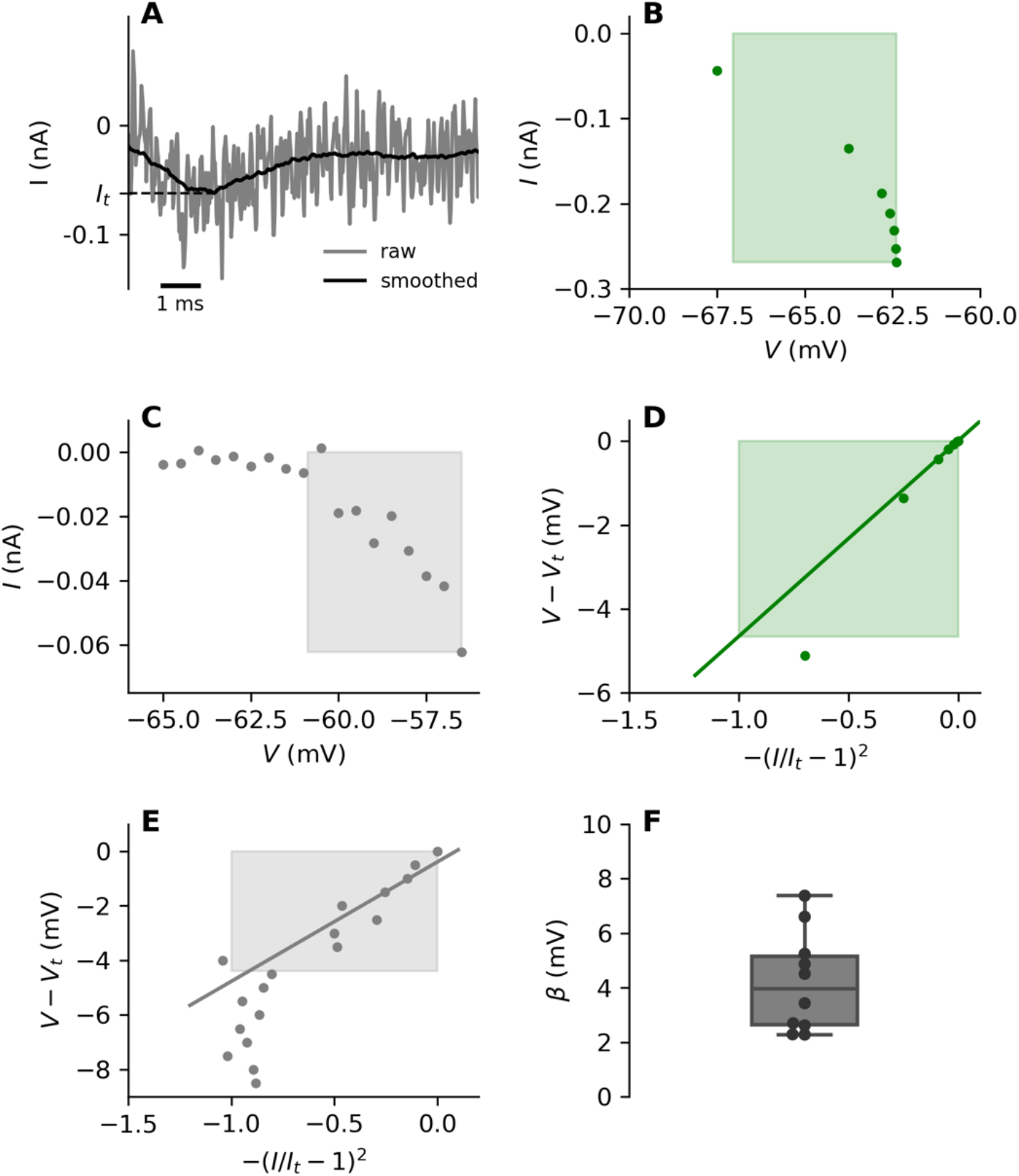
Axial current near threshold. A, Current recorded at threshold (gray). After smoothing (black), It is the peak value. B, Peak current vs. V in a biophysical model with extended AIS. Threshold was measured by bisection for better precision. The shaded box represents the region 3 mV below threshold. C, Peak current vs. V measured below threshold in a RGC. D, Difference between membrane potential and voltage threshold vs. quadratic normalized current in the biophysical model, with the regression line (slope β = 4.7 mV). E, Same as D in the RGC (slope of regression line: β = 4.4 mV). F, Slope of linear regressions shown in E over all cells.

#### Threshold vs. AIS geometry

Both the voltage and axial current at threshold are predicted to depend on AIS geometry, namely to decrease when the AIS is shifted away from the soma, all else being equal. We analyzed these relations in *n* = 10 cells (cells were excluded either because AIS geometry was not measured or reference potential drifted). There was no significant linear correlation in our data between voltage threshold and either AIS start position (Fig. 6A, *p* = 0.54, Pearson test) or length (Fig. 6B, *p* = 0.14, Pearson test). However, voltage threshold varies theoretically with both quantities as −*k* log(*x*_1/2_*L*). The correlation was stronger with log(*x*(*x*_1/2_*L*), although still weak (Pearson correlation r = 0.62, *p* = 0.06). The regression slope was *k* = 4.3 mV, a plausible value (Fig. 6C). We note that diameter and perhaps conductance density, which both contribute to the voltage threshold, may also vary across cells.

**Figure 6.**
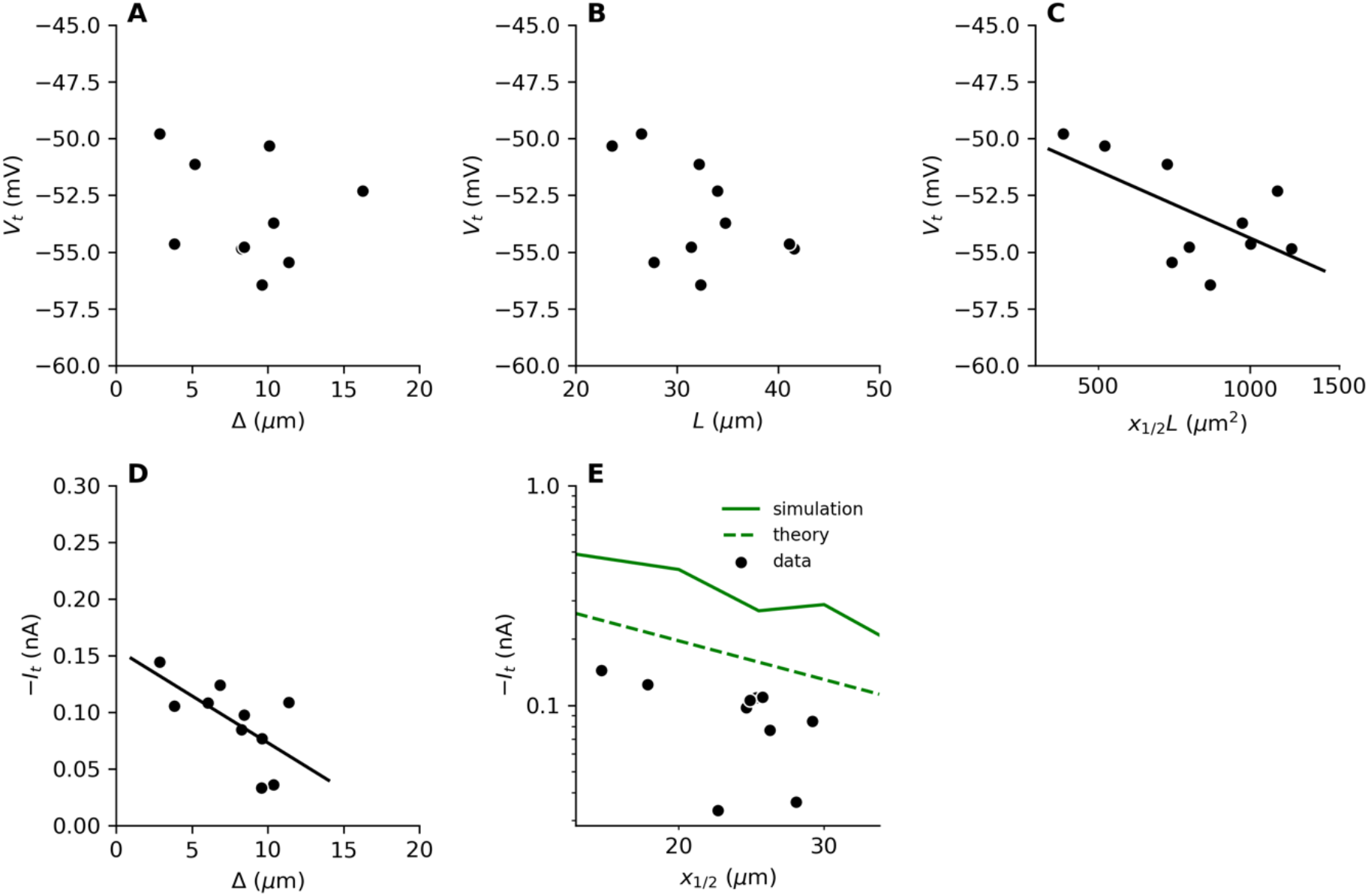
Threshold vs. AIS geometry. A, Voltage threshold vs. AIS position Δ is RGCs. B, Voltage threshold vs. AIS length L. C, Voltage threshold vs x_1/2_L in logarithmic space, with logarithmic regression line (slope k = 4.3 mV; p = 0.06, Pearson test). D, Current at threshold vs. AIS position, with regression line (p = 0.04, Pearson test). E, Current at threshold vs. AIS middle position x1/2, with theoretical prediction using d = 1 μm (dashed green) and simulation in a biophysical model (solid).

We observed an inverse correlation between axial current threshold and AIS position (Fig. 6D, *p* = 0.04, Pearson test). Theory makes a quantitative prediction: *I*_*t*_ = *k*/*R_a_*, with *R_a_* measured from the soma to the middle of the AIS. This may differ by a constant factor in a complex biophysical model (Fig. 6E, compare dashed and solid lines). Measured currents are lower than predicted and the inverse correlation is barely significant (Pearson correlation r = −0.65, *p* = 0.08). As explained above, underestimation and limited precision were expected. Nonetheless, the magnitude of measured currents was reasonably close to theoretical estimations (92 pA vs. 160 pA on average, with all but two cells between 70 and 150 pA).

### Adaptation of the axial current

#### Properties of adaptation

We observed that the axial current at spike initiation has just the right magnitude to depolarize the soma to the somatic regeneration threshold. What would happen if the availability of sodium channels varied? In many neurons, sodium channels can inactivate substantially below threshold, producing voltage threshold adaptation (Azouz & Gray, 2000; Platkiewicz & Brette, 2011; Fontaine *et al.*, 2014). Threshold adaptation has been observed in current-clamp recordings of salamander RGCs (Mitra & Miller, 2007). If this phenomenon reflects the inactivation of AIS sodium channels, it may compromise the transmission of the AIS spike to the soma.

We examined this issue by holding the neuron at different potentials *V_0_* before measuring the voltage threshold. We observed that the threshold increases substantially with *V_0_*(Fig. 7A). The relation between *V_t_* and *V_0_* follows the theoretical expectation for threshold adaptation due to sodium channel inactivation (Platkiewicz & Brette, 2011; Fontaine *et al.*, 2014), where the threshold starts increasing above the baseline *V_min_* when *V_0_* exceeds the half-inactivation voltage *V_i_* of sodium channels, with a slope *k*_*i*_/*k*_*a*_ ≈ 1 in the depolarized range (where *k_i_* and *k_a_* are the inactivation and activation slope factors, respectively). By fitting the theoretical relation, we find *V*_*i*_ ≈ −55.8 ± 3.1 mV (Fig. 7B), *V*_*i*_ − *V*_*min*_ ≈ −0.7 ± 2.9 mV (Fig. 7C), *k*_*a*_ ≈ 4.1 ± 2.2 mV (Fig. 7D), and *k*_*i*_/*k*_*a*_ ≈ 0.9 ± 0.18 (Fig. 7E). These values are consistent with expectations if threshold adaptation is due to sodium channel inactivation.

**Figure 7.**
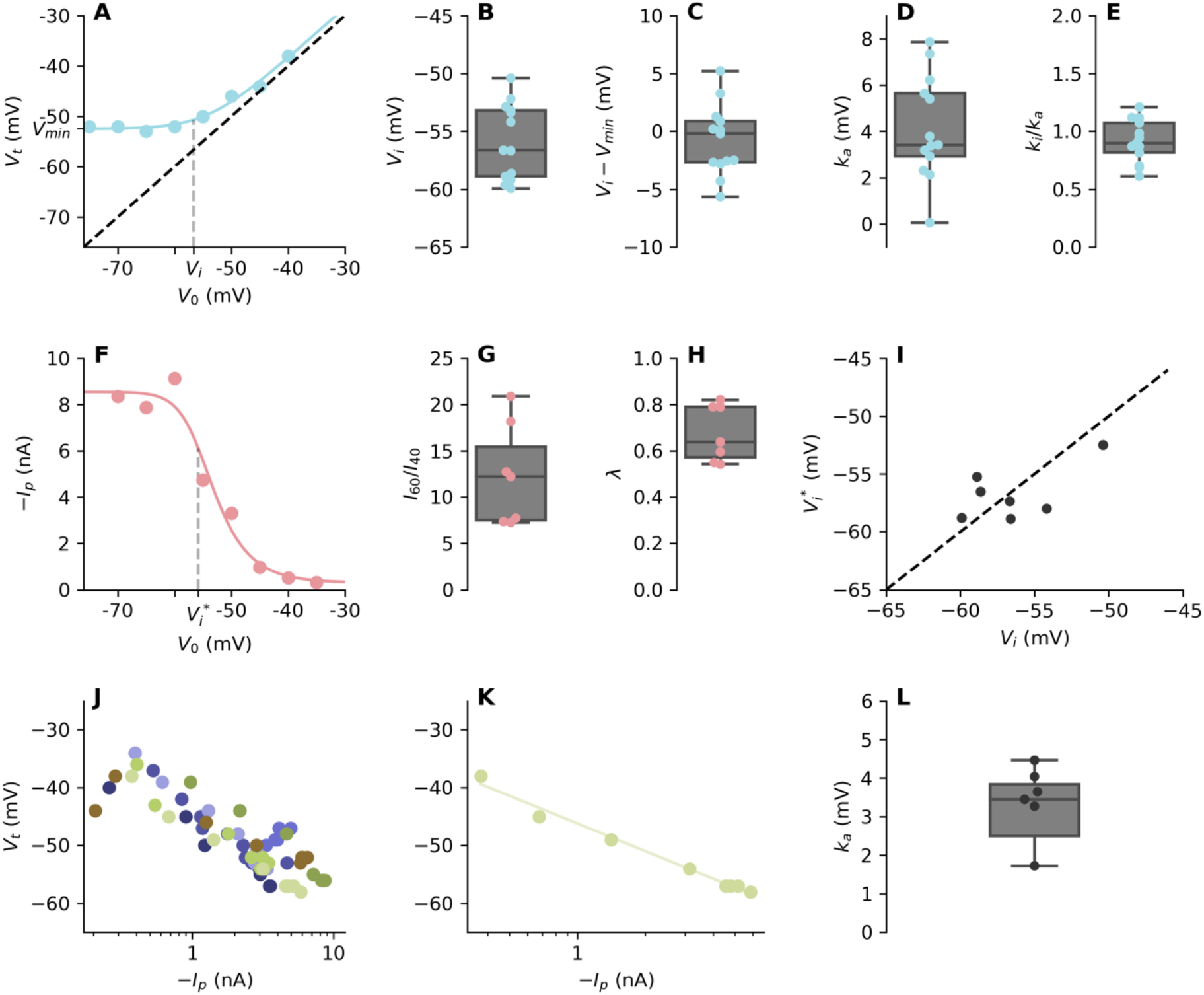
Adaptation of the axial current at spike initiation. A, Voltage threshold vs. initial holding potential V0 in a RGC. The dashed line is the identity V_t_ = V_0_, and the solid curve is a fit to the theoretical relation. B, Statistics of half-inactivation voltage Vi from theoretical fits. C, Statistics of Vi-Vmin from fits. D, Statistics of activation slope k_a_. E, Statistics of k_i_/k_a_. F, Axial current Ip at spike initiation vs. initial holding potential V_0_ in a RGC. The dashed line shows the potential 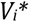 where Ip is attenuated by √2. G, Current attenuation I_60_/I_40_ from −60 to −40 mV, over all cells. H, Current at half-inactivation voltage Vi, relative to the maximum current. I, Comparison between 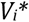 obtained from current attenuation and Vi obtained from voltage threshold adaptation (the diagonal line is the identity). J, Voltage threshold Vt vs. axial current in logarithmic space, over all RGCs (n = 9; each color corresponds to one cell). K, Voltage threshold Vt vs. axial current for one RGC, with logarithmic regression line (half-slope k = 3.4 mV, r = 0.99). L, Statistics of ka from logarithmic regressions over all cells.

We then measured the axial current at spike initiation (just above threshold) as a function of *V_0_*(note that there are fewer data points because current recordings were discarded when *R_s_* changed by more than 30%). We observed that the current decreased considerably with increasing *V_0_*(Fig. 7F). On average, it attenuates by a factor 12.3 ± 5.1 when *V_0_*increases from −60 to −40 mV (Fig. 7G). At *V_i_*, the current is 32 ± 10 % smaller than the maximum current (Fig. 7H).

If adaptation of voltage threshold and axial current are both due to sodium channel inactivation, then axial current and voltage threshold should co-vary with V_0_. Theoretically, *V_t_* varies with available conductance *g* as −*k* log *g* (Platkiewicz & Brette, 2011). For low *g*, the axial current *I_p_* is proportional to √*g*. Therefore, *V_t_* should vary with *I_p_* as 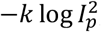, or equivalently, −2*k* log |*I_p_*|.

We first note that the potential 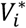 at which the axial current is attenuated by √2 is indeed close to the half-inactivation voltage *V_i_* estimated from threshold adaptation 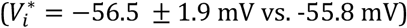 (Fig. 7I). Then when we compare *V_t_* with *I_p_*, we find a logarithmic relation (Fig. 7J, K) with half-slope *k*_*a*_ = 2.4 ± 2.7 mV (Fig. 7L). Note that this average includes one outlier; the median k_a_ is 3.4 mV (the smaller number of points is due to the fact that exclusion criteria for both *V_t_* and *I_p_* are applied). This strongly suggests that both threshold and axial current adaptation are due to the same phenomenon, sodium channel inactivation.

While the axial current above threshold is strongly modulated by the available Na^+^ conductance, resistive coupling theory predicts that the threshold axial current depends on AIS geometry but not on sodium conductance (*I_t_* = *k/R_a_*). Figure 8A shows current-voltage relations for different V_0_ in the same cell. The curves appear to shift horizontally when V_0_ is changed, so that the voltage threshold increases with *V_0_*but the axial current at threshold does not, as shown specifically on Figure 8B. Over all measured cells (*n* = 6; voltage threshold and current threshold were only measurable with a stable *R_s_* in a few cells), *I_t_* varied by a factor smaller than 2.5 (1.2 ± 0.6) between −60 and −40 mV (Fig. 8C), whereas *I_p_* varied by a factor 12.3 on average.

**Figure 8.**
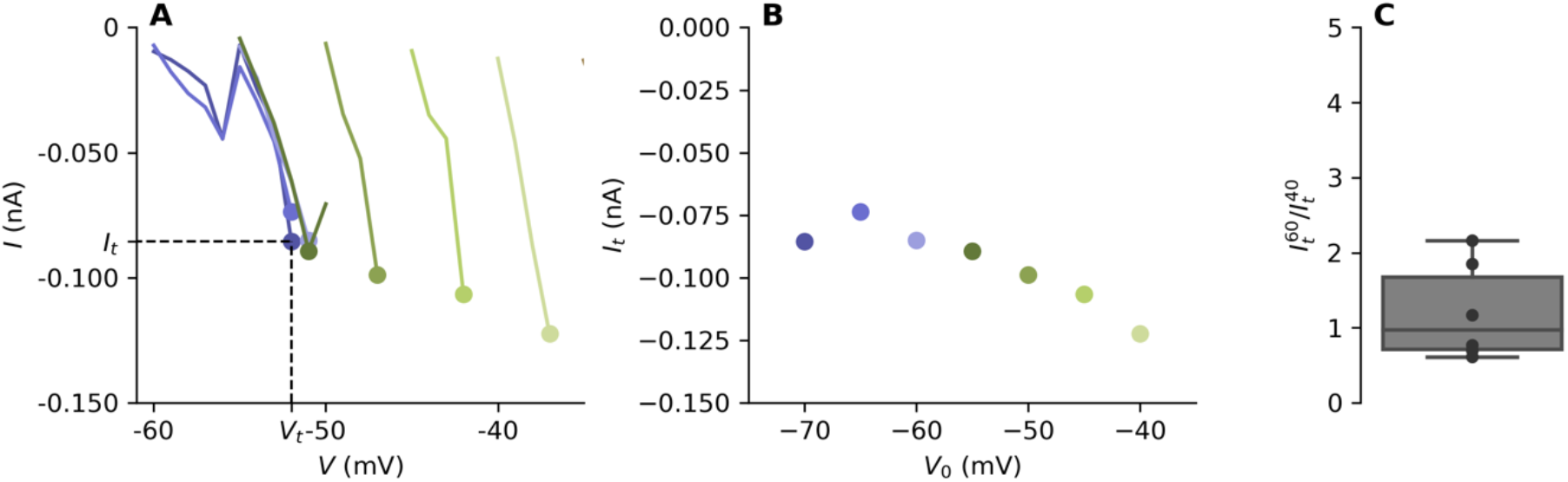
Adaptation of the axial current at threshold. A, Current vs. somatic potential V a few mV below threshold, shown for different initial holding potentials V0 (−70, dark purple to −40 mV, light green) in the same RGC. B, Current at threshold vs. V0 in the same cell. C, Attenuation of threshold current 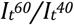.

#### Compensation of axial current attenuation

Figure 9A shows the attenuation of *I_p_* as a function of *V_0_*in one cell. Here *I_p_* attenuates by a factor 7.3 between −60 and −40 mV. If the axial current at spike initiation attenuates by a factor 7, then we expect the induced somatic depolarization to also attenuate by a factor 7, to about 4 mV, which seems insufficient to reach the threshold for somatic regeneration. However, this is not what we found. In this cell, the total transmitted charge, obtained by integrating the current, attenuates only by a factor 1.7 (Fig. 9B). This occurs because current duration increases at high *V_0_*(Fig. 9C). Over all measured cells (*n* = 7), transmitted charge attenuated by a factor 3.1 ± 1.4 from −60 to −40 mV, compared to 12.3 ± 5.1 for the axial current (Fig. 9D). The increase in current duration was observed consistently above −50 mV (Fig. 9E).

**Figure 9.**
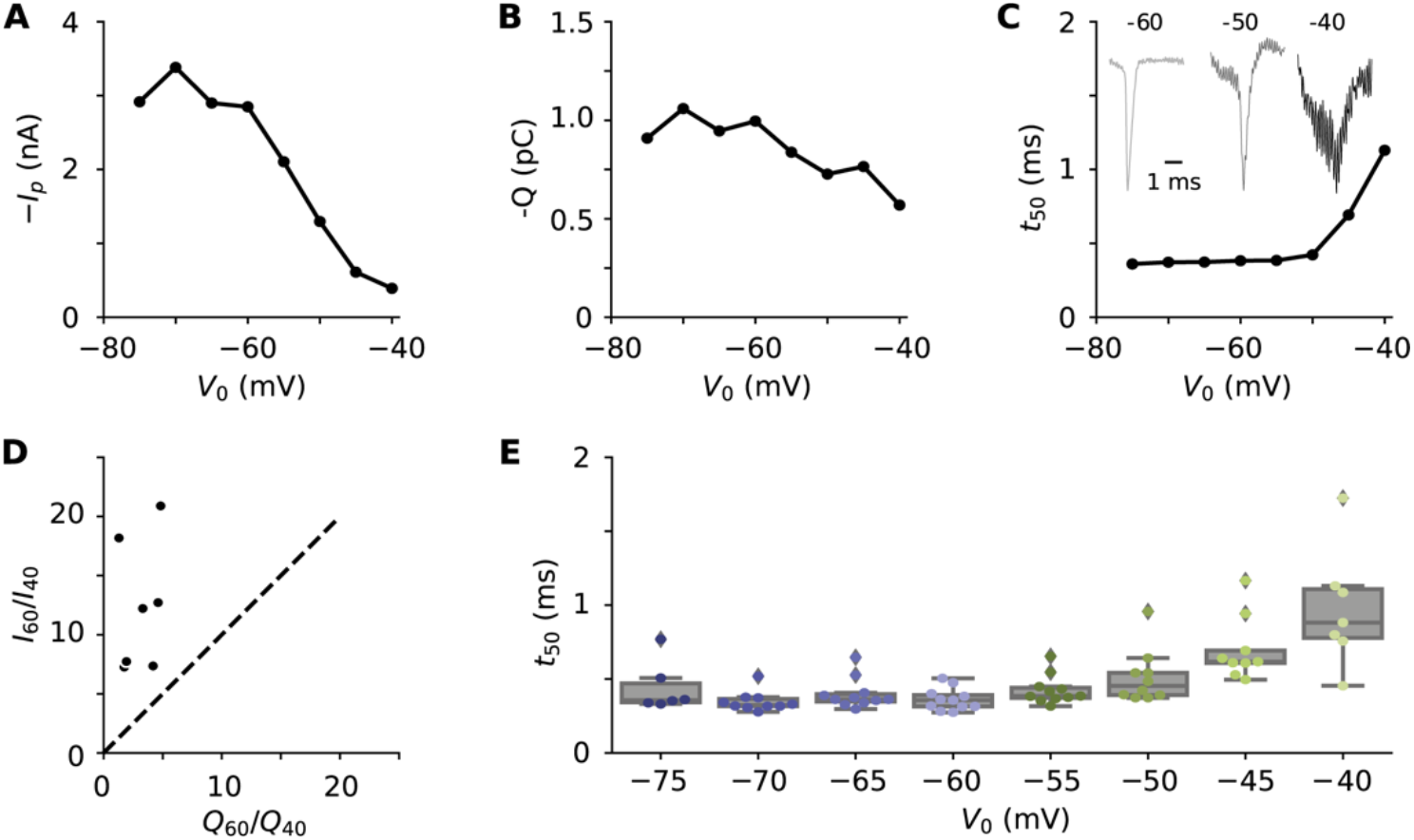
Compensation of axial current attenuation. A, Axial current at spike initiation vs. initial potential V_0_ in a RGC. B, Total transmitted charge Q vs. V_0_ in the same cell. C, Current duration t_50_ vs. V_0_ in the same cell. Top traces: normalized axial currents measured at V_0_ = −60 mV (light gray), −50 mV (dark gray) and −40 mV (black). D, Current attenuation vs. charge attenuation between −60 and −40 mV, over all cells. E, Current duration t_50_ vs. V_0_ in all cells.

Indeed, we could occasionally observe spontaneous bursts on top of a depolarizing wave, with APs triggered at potentials up to −40 mV, with no sign of transmission failure. An example is shown in Figure 10A, with a selection of individual APs shown in Figure 10B, and corresponding phase plots in Figure 10C. During this burst, spike onset increased up to about −40 mV while the somatic regeneration threshold was stable (Fig. 10D).

**Figure 10.**
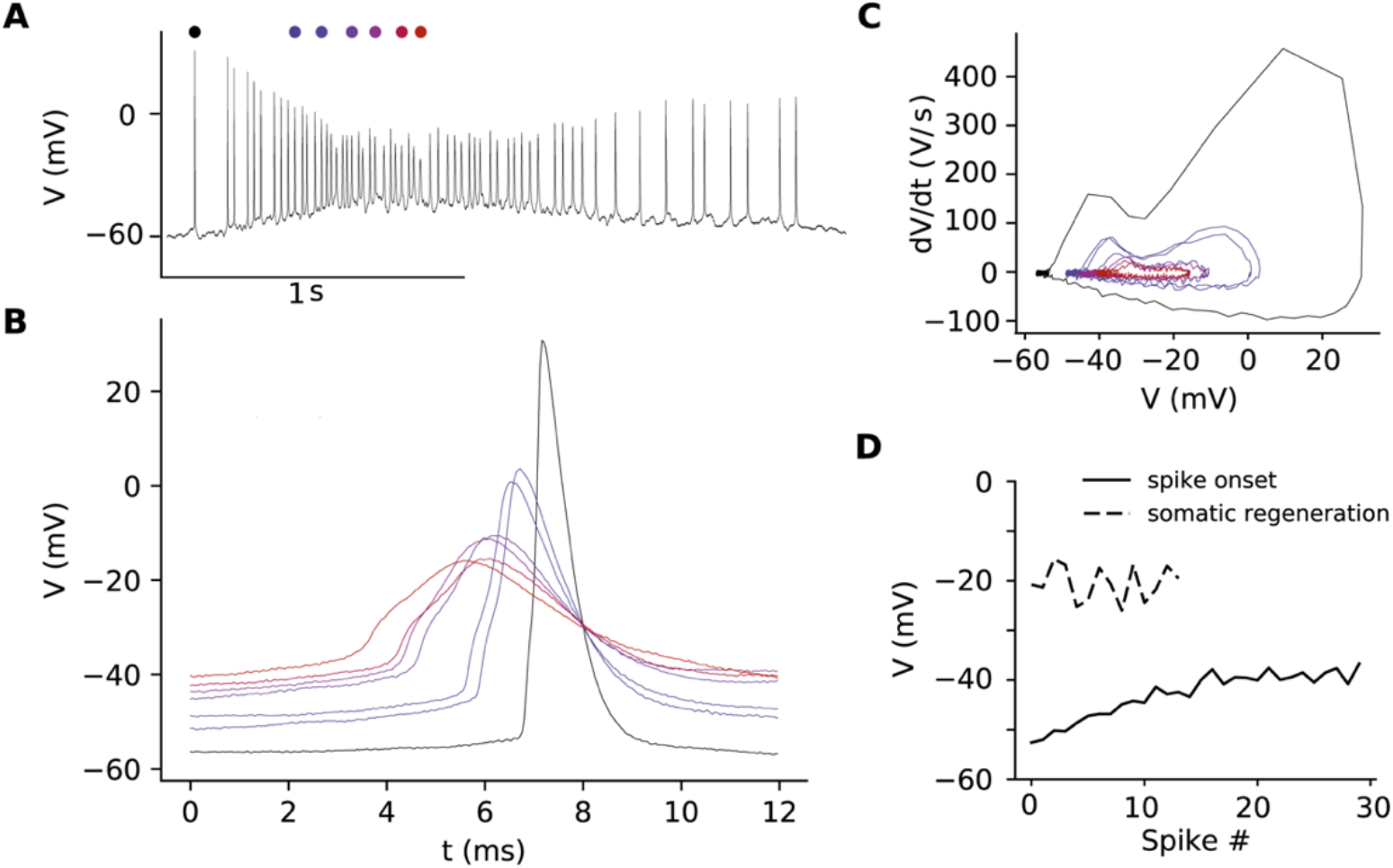
Changes in AP properties during a spontaneous burst. A, Spontaneous burst of APs in a RGC. B, Individual APs marked in A. C, Phase plot of Aps marked in A. D, Spike onset and somatic regeneration threshold of the successive APs in A. Somatic regeneration threshold could only be accurately measured for the first 12 spikes because of noise.

Thus, detailed properties of the axial current appear to be such as to ensure reliable AP transmission to the soma in changing conditions.

## Discussion

### Summary

In summary, we have observed that the AIS of RGCs produces a large axial current at spike initiation (about 7 nA), which requires a high Na^+^ conductance density (most likely several thousand S/m^2^). The charge that this current transmits to the soma co-varies with somatic capacitance, in such a way as to produce a depolarization of about 30 mV, the amount necessary to bring the somatic potential to spike regeneration threshold. Theory shows that the axial current is mainly determined by AIS position and diameter, and to some extent by Na^+^ conductance density, but perhaps counter-intuitively not by AIS length.

In agreement with resistive coupling theory (Brette, 2013; Kole & Brette, 2018), the axial current is small below threshold (on the order of 100 pA at threshold, and undetectable a few mV below) and decreases when the AIS is further away from the soma, which reduces energy consumption.

We have also observed that the voltage threshold for spike initiation adapts to depolarization, in a way compatible with Na^+^ channel inactivation. Consistently, the axial current at spike initiation also decreases when the threshold adapts. This attenuation can reach a factor of 10 or more for large depolarizations, which could potentially compromise spike transmission to the soma. However, we found that this attenuation is compensated by a broadening of the axial current.

Overall, our results are in good agreement with predictions of resistive coupling theory. The inferred Na^+^ channel activation slope factor, which was found consistently to be *k* ≈ 4 mV in several distinct data sets, may seem to be on the low end of Boltzmann fits to patch-clamp recordings, typically 4-8 mV (Angelino & Brenner, 2007). However, this is likely because this parameter is generally obtained from fits on a broad voltage range, while an exponential fit around the spike initiation voltage yields lower values (Platkiewicz & Brette, 2010). For example, Hodgkin and Huxley (1952) found that the Na^+^ current-voltage curve of the squid axon was well fitted by an exponential of slope 4 mV; Baranauskas and Martina (2006) also noted that in cortical pyramidal cells, the slope was lower when estimated around spike threshold than on a broader range (5.4 mV vs. 6.4 mV).

### Limitations

One of the main technical limitations to interpret the results of this study is that axonal diameter *d* cannot be measured precisely with conventional optical microscopy. This is an important limitation because theory shows that key properties are very sensitive to diameter. Specifically, axial resistance is inversely proportional to *d^2^*. This results in an error in resistance estimation of around 50% for a 200 nm error in axon diameter estimation (assuming *d* ≈ 1 μm). This translates to comparable errors in axial current predictions. This limitation should also be kept in mind when interpreting other studies where changes in AIS geometry are observed (see below). The best way to overcome this limitation would be to measure axonal diameter precisely using either electron microscopy or super-resolution microscopy.

Two other limitations are the measurement of intracellular resistivity R_i_ and axonal tapering. Axial resistance is proportional to intracellular resistivity *R_i_*, but this parameter is difficult to estimate. Ideally, it should be measured by simultaneous recordings in the axon and soma, and a precise estimate requires a precise measurement of axon diameter. Stuart and Spruston (1998) estimated *R_i_* = 70-100 Ω.cm in dendrites of cortical pyramidal cells, based on simultaneous recordings in soma and apical dendrite. Although this value is mainly determined by the concentration of the most mobile ions (i.e., mainly K^+^), which is not expected to vary widely across the cell, it is conceivable that it is higher in thin crowded structures such as the proximal axon. Higher values, up to 150 Ω.cm, have been used in modeling studies of RGCs (Sheasby & Fohlmeister, 1999; Fohlmeister *et al.*, 2010), but these are based on model optimization using somatic recordings.

In the theory, we did not take into account axonal tapering. However, in RGCs, axon diameter decreases from soma to AIS, then decreases again along the AIS (Raghuram *et al.*, 2019). The theory assumes a uniform diameter, because general analytical solutions do not exist with variable diameter. For the calculation of the axial current at spike initiation, taking into account tapering would tend to reduce the axial resistance between soma and AIS, as if the AIS were closer to the soma. The diameter in the calculation of δ should be the diameter of the proximal AIS.

We were not able to confirm the inverse relation between AIS position and axial current at spike initiation that theory predicts. A plausible reason is that the axial current is very sensitive to diameter, which might have varied substantially across cells. It is also possible that the available Na^+^ conductance density varied across cells. Indeed, our data on adaptation show that half-inactivation voltage (about − 57 mV) is close to the initial potential used in our voltage clamp measurements of axial current (−60 mV). Another potential source of variability is that we have studied mice at an age where there are developmental changes in the expression of Na^+^ channel subtypes (Boiko *et al.*, 2003; Van Wart *et al.*, 2007), which might have contributed some variability. Finally, series resistance introduces errors that may have been incompletely corrected offline. This issue could be addressed by measuring currents with two electrode voltage-clamp (Barrett & Crill, 1980).

### Na^+^ conductance density

Whether the AIS has high Na^+^ conductance density has been controversial (Colbert & Pan, 2002; Kole *et al.*, 2008; Fleidervish *et al.*, 2010). This question has been typically considered from the viewpoint of excitability: high conductance density has been proposed to account for the fact that the AIS spike initiates about 30 mV below the somatic regeneration threshold. Other contributing factors are the lower activation threshold of axonal sodium channels (Kole & Stuart, 2008; Hu *et al.*, 2009) and the axial resistance between soma and AIS (Brette, 2013). Here we examined another empirical constraint on Na^+^ conductance density, the axial current that the AIS generates at spike initiation.

By just considering the area of the AIS, to produce a current of 6.7 nA requires a conductance density of about 1150 S/m^2^ with *d* = 0.7 μm (average diameter across the AIS in our data) or 800 S/m^2^ with an upper estimate of *d* = 1 μm. This is a lower bound that neglects considerations of cable theory, namely the fact that the axial current flows from the distal end of the AIS to the soma.

It is possible to calculate the maximum axial current produced by an axon of diameter *d* and conductance density *g*. This calculation shows that, to account for a current of 6.7 nA, *g* must be at least 1200 S/m^2^ if *d* = 1 μm, and about 2500 S/m^2^ if *d* = 0.8 μm, independently of AIS position. Here the relevant diameter is the diameter of the proximal AIS (about 0.9 μm in our data). Finally, taking into account measured AIS position, the data are consistent with *g* around 5000 S/m^2^ (with *d* = 1 μm), although the minimum is broad. Overall, this analysis indicates that *g* should be several thousand S/m^2^. This estimate is independent of Na^+^ channel kinetics, and in particular it holds even if Na^+^ channels cooperate (Naundorf *et al.*, 2006).

Lorincz and Nusser (2010) counted 187 Nav1.6 channels per μm^2^ in the AIS of CA1 pyramidal cells. Assuming a unitary conductance of 10-20 pS per channel (Hille, 2001), this amounts to 1870-3740 S/m^2^. However, this is an estimate of the structural density, not necessarily of the functional density. In a computational model of layer 5 pyramidal cells, a density of 2500 S/m^2^ was necessary to account for the measured initial depolarization speed of somatic APs (Kole *et al.*, 2008). Similarly, optimization of a model of RGCs for AP shape yielded a conductance density of about 5000 S/m^2^ (Guo *et al.*, 2013). Our analysis provides an estimation that is less dependent on model specifics, and confirms these previous studies.

The theoretical analysis indicates that a high conductance density is likely a necessary condition to transmit the AIS spike to the soma in a variety of cell types, due to the drastic geometrical variation at the axosomatic boundary. The minimum conductance density to produce an axial current *I* is proportional to I^2^/d_AIS^3^_. If we assume that the current must scale with the area of the soma, then the minimum *g* is proportional to d_soma^4^_/d_AIS^3^_. This ratio appears to be approximately conserved across cell types (Goethals & Brette, 2020), and therefore most neurons should face the same constraint requiring a similar conductance density in the AIS.

It should be noted that, despite a high conductance density at the AIS, the total Na^+^ influx through the AIS should theoretically have the same order of magnitude as through the soma and proximal dendrites, as observed (Fleidervish *et al.*, 2010). Indeed, the total Na^+^ influx at the AIS should match the charge necessary to depolarize the soma by about 30 mV, while the total Na^+^ influx at the soma (and proximal dendrites) should account for a further depolarization of a few tens of mV (about 45 mV in our cells). The AIS influx should occur preferentially in the proximal AIS, even if conductance density is uniform, because the driving force of the Na^+^ channel is larger there (see Fig. 4A). This has indeed been observed in cortical pyramidal cells (Baranauskas *et al.*, 2013).

### Structural tuning of the AIS

In layer 5 cortical pyramidal cells, Hamada et al. (2016) observed that AIS position was inversely related to the diameter of the apical dendrite. Quantitatively, this relation was consistent with a proportionality relation between the axial current produced by the AIS and the somatodendritic capacitance. Here we showed more directly that, in RGCs, the charge transmitted by the AIS covaries with the somatodendritic capacitance, in such a way as to depolarize the soma to the threshold for somatic spike regeneration.

Overall, our measurements are in line with quantitative predictions of resistive coupling theory. However, we did not observe a correlation between AIS position and capacitance. Theoretically, the structural parameters that determine the axial current are AIS position and diameter (and not AIS length, at least not in the range of observed lengths). Therefore, one would expect a negative correlation between AIS position and capacitance if diameter were homogeneous across cells, or at least uncorrelated to capacitance. In fact, Raghuram et al. (2019) observed a positive correlation between AIS position and soma size in *α* S RGCs. Such a correlation would be expected if AIS diameter scaled with soma size. The authors did observe a positive correlation between soma size and the diameter of the proximal axon (we note that observing such correlations for the AIS proper, which is below 1 μm in diameter, may not be feasible). It cannot be excluded that the lack of significant inverse correlation between AIS position and capacitance is due to the limited precision of our measurements, especially as we did observe an inverse correlation between AIS position and threshold axial current. It is possible that the availability of Na^+^ channels varied across cells. In any case, we stress that axon diameter is a key structural parameter in setting the axial current as well as excitability, and therefore it must be considered to correctly interpret experimental results.

It remains that, in order to produce an axial current of appropriate magnitude from an AIS of a given diameter and conductance density, the AIS must be positioned appropriately. A number of studies have shown that AIS position can vary across cells (Kuba *et al.*, 2006; Hamada *et al.*, 2016; Höfflin *et al.*, 2017), during development (Galiano *et al.*, 2012; Kuba *et al.*, 2014; Gutzmann *et al.*, 2014), and with activity (Kuba *et al.*, 2010; Grubb & Burrone, 2010; Grubb *et al.*, 2011; Kuba, 2012; Evans *et al.*, 2015; Jamann *et al.*, 2017, 2020). These changes have often been suggested to reflect a homeostatic regulation of excitability, but theory predicts a small effect of AIS position on excitability (Goethals & Brette, 2020), and a large effect on axial current. Therefore, it is conceivable that these changes reflect a homeostatic regulation not of excitability *per se*, but of the axial current required to transmit the AIS spike to the soma. For example, Grubb and Burrone (2010) report that when cultured hippocampal neurons are depolarized with 15 mM KCl, capacitance decreases by about 10% while the AIS shifts away from the soma. This distal shift is consistent a decrease in axial current required to match the decrease in capacitance. In the same way, developmental changes in AIS position may also be consecutive of changes in somatic diameter or dendritic area. Finally, in neurons where backpropagation of the AP to the soma may be undesirable, such as some types of auditory neurons (Kuba *et al.*, 2006; Scott *et al.*, 2007), a distal placement of the AIS may be beneficial.

### Adaptation

Threshold adaptation has been observed in current-clamp recordings of salamander RGCs (Mitra & Miller, 2007), as well as in many other cell types (reviewed in (Platkiewicz & Brette, 2011)). We also observed it in mouse RGCs and quantified it precisely. The voltage threshold starts increasing when the membrane is depolarized above ≈ −56 mV, and for large depolarizations the slope of the relation between potential and threshold is close to 1. That is, the threshold tracks the membrane potential so as to remain a few mV above it. These observations are consistent with theoretical expectations based on Na^+^ channel inactivation (Platkiewicz & Brette, 2011). In layer 5 pyramidal cells, half-inactivation voltage of AIS Na^+^ channels is about −61 mV (Kole & Stuart, 2008), which is in line with our observations.

To our knowledge, adaptation of the axial current had not been reported before. Our quantitative analysis shows that the co-variation of axial current and threshold is consistent with AIS Na^+^ channel inactivation being the cause of both phenomena. The axial current attenuated by a factor 12 on average over a 20 mV depolarization. This would reduce the charge transmitted to the soma by the same factor and possibly compromise spike transmission to the soma, if the current spike shape were unchanged. However, we observed that this attenuation was largely compensated by a broadening of axial currents. This means that the AP at the AIS broadens when the soma is depolarized. In fact, such broadening has been observed in layer 5 pyramidal cells and attributed to the inactivation of Kv1 channels (Kole *et al.*, 2007). As Kv1.2 is expressed in the distal AIS of RGCs (Van Wart *et al.*, 2007), this might explain our observations.

In conclusion, our observations indicate that structural and channel properties of the AIS are functionally organized in such a way as to ensure reliable transmission of the AP to the soma.

## Materials and Methods

All data are available at https://zenodo.org/record/4005629#.X00H9y3pP-Z. Code for both data analysis and model simulations is available at https://github.com/romainbrette/AIS-geometry-and-axial-current.

### Ethical statement

Timed-pregnant Swiss and C57BL/6NRj mice were purchased from Janvier Labs (Le Genest Saint Isle, France) and housed under controlled conditions (22 ± 1°C, 60 ± 10% relative humidity, 12/12h light/dark cycle, food and water *ad libitum*). All animal procedures were performed in strict accordance with institutional guidelines and approved by local ethics committees (C2EA-05: Comité d’éthique en expérimentation animale Charles Darwin) and by European Communities Council Directive 2010/63/UE.

### Whole-cell electrophysiology of RGCs

Mice were taken at postnatal day 10-12 (P10-12). The pup was rapidly decapitated, the eyes were removed and placed in Ringer’s medium containing (in mM): 119 NaCl, 2.5 KCl, 1.0 KH_2_PO_4_, 11 glucose, 26.2 NaHCO_3_, 2 CaCl_2_ and 1 MgCl2 (290-295 mOsm), bubbled with carbogen (95% O_2_/5% CO_2_). The retina was dissected and fixed on filter paper over a small hole (N8895, Sigma-Aldrich) with the RGC layer upwards and continuously perfused with Ringer’s solution warmed to 32 degrees Celsius.

Thick-walled borosilicate pipettes (OD/ID of 1.5/0.87 mm; 30-0060, Harvard Apparatus) were pulled on a P-1000 Flaming/Brown puller (Sutter Instruments). Pipettes were filled with intracellular solution containing (in mM): 128 K-gluconate, 10 HEPES, 16 KCl, 1 EGTA, 2 Mg-ATP, 0.5 Na_2_-GTP, pH 7.25 with KOH (275 mOsm) and 1-2 mg/mL biocytin (B4261, Sigma-Aldrich). Open tip resistance was 2.5-4 MΩ. Reported potentials were corrected for a liquid junction potential of −11 mV. Whole-cell recordings were made with a Multiclamp 700B amplifier (Axon Instruments), filtered at 10 kHz and digitized at 50 kHz using a DigiData 1440A (Axon Instruments) and Clampex 10.7 running on Windows 10. High-resistance patch seals (>1 GΩ) were obtained before breaking into the cell. Recordings with a series resistance *R_s_* above 25 MΩ, or with a residual *R_s_* (after compensation) above 5 MΩ, were discarded. The resting membrane potential of the cell was recorded in the first minute after breaking in.

Passive cell properties were recorded by stepping from −70 to −80 mV in voltage-clamp mode without whole-cell compensation. Series resistance was electronically compensated 80-95% with a lag of 18 μs. Between protocols we repeated the voltage step without compensation to monitor changes in series resistance, and series resistance compensation was adjusted if necessary. Passive currents were subtracted using a P/n protocol (5 steps of 5 mV) that preceded each protocol. The P/n protocol was missing for a few recordings; we then subtracted the passive response using a 10 mV step.

Adaptation protocols started with a long adaptation step at *V_0_*(0.5 s, *V_0_* varied by steps of 5 mV) followed by an activation step (resolution of 1 mV) to elicit an AIS spike. We ensured that the adaptation step was long enough by varying the step duration in a few cells.

In current-clamp mode, bridge balance and pipette capacitance cancellation (6.2-7.1 pF) were used. Hyperpolarizing current pulses were injected to measure the cell’s capacitance (see *Electrophysiological data analysis*). Next, for n = 16 cells, 5-20 minutes of spontaneous activity were recorded to analyze spontaneous APs.

At the end of the experiment, the pipette was retracted to obtain an outside-out patch. Outside the retina the tip was cleaned with brief, positive pressure to remove the remaining membrane patch and the potential offset was noted to check for any drift in the reference potential.

The retina was rinsed in 0.12 M PBS, fixed for 15 minutes in 4% paraformaldehyde in 0.12 M phosphate buffer, rinsed again 2 times with 0.12 M PBS and kept in 0.12 M PBS at 4 degrees Celsius for immunolabeling procedure at a later stage.

### Immunohistochemistry

A few days after recording, retinas were washed during 5 minutes in PBS and permeabilized overnight at 4°C in 5% normal goat serum (NGS, Sigma Aldrich) and 1% Triton-X100 in PBS. Retinas were then incubated with a solution of 0.1M PBS containing the mouse monoclonal anti-ankyrin-G (clone N106/36; 1:250; NeuroMab; ID: AB_2749806) and 1% normal donkey serum (NDS) for 24h at 4°C. Retinas were washed 4 x 15 minutes in 0.1M PBS and then incubated with a PBS solution containing donkey anti-mouse Alexa Fluor 488 (1:500, Thermo Fisher; ID: AB_141607), Streptavidin Alexa Fluor 594 (1:500, Thermo Fisher, ref. S11227), 5% NGS and 0.1% Triton-X100 overnight at 4°C. Retinas were washed 4×15 minutes in PBS and mounted onto SuperfrostΩ slides and coverslipped with Fluoromount. Mounted retinas were stored at 4°C until confocal imaging.

### Confocal imaging

The retinas labelled for biocytin and ankyrin-G were sequentially captured with an inverted laser scanning confocal microscope (FV1000, Olympus) equipped with an Argon (488nm) ion laser and laser diode (555 nm), filter cubes appropriate for Alexa Fluor 488 and Alexa Fluor 594, and a 40x objective (oil, NA 1.3). Acquisition settings were optimized for each cell. Z-stacks were obtained with a step size of 0.5 μm. Confocal x-y sampling resolution was 0.155 μm, except for three cells for which it was 0.207 μm.

### Electrophysiological data analysis

Data were analyzed with custom Python scripts.

#### Estimation of passive properties

The raw series resistance 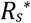 was measured from responses to a test pulse in voltage clamp: *R*_*s*_ = Δ*V*/*I*_0_, where ΔV is the voltage pulse amplitude and *I_0_* is the amplitude of the first transient peak. The residual series resistance *R*_*s*_ during a given recording is 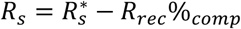 where R_rec_ is the series resistance used for compensation during the experiment and %comp is the amount of compensation. Effective capacitance is estimated from the response to current pulses, by fitting an exponential to the first ms, which is the time scale of the axial current. This estimation was done in n = 17 cells.

#### Analysis of APs

The first AP recorded during spontaneous activity was used to measure AP features (Fig. 1). Spontaneous activity was recorded in 16 cells; 6 of them were excluded from this analysis because the reference potential drifted by more than 3 mV. To compute the phase plots (*dV/dt* vs *V*), we ensured that the plotted points are isochronic, by considering that *dV/dt* corresponds to the derivative midway between two consecutive points, and interpolating the values of *V* at that midpoint. Spike onset was defined as the potential when *dV/dt* crosses 20 mV/ms for the last time before the AP peak. The value of *dV/dt* for the initial segment component was defined as the first local maximum between spike onset and the global maximum of *dV/dt*. In a few cells, this was equal to the global maximum. The regeneration threshold is defined as the potential at the point of maximal acceleration *d^2^V/dt^2^* after the initial segment component.

#### Correction of series resistance error

Axial currents were corrected using a minor adjustment of the method described by Traynelis (1998). The presence of the series resistance results in an error in clamping the somatic potential equal to - *R_s_.I_e_*, where *I*_e_ is the current through the electrode. This produces a capacitive current through the somatic membrane equal to *C.dV/dt* = -*R_s_C dI_e_/dt*, which results in filtering the axial current through a low-pass filter with time constant *τ* = *R*_*s*_*C*s. We correct the recorded current by subtracting this capacitive current:

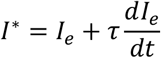

where I* is the corrected current. The time constant is estimated directly by fitting an exponential to the first 0.5 ms of the response to a small voltage step (with the same amplifier tunings as for subsequent recordings) (Fig. 11A). Note that as the whole cell compensation circuit of the amplifier was used, this must be understood as a residual time constant, which can be negative. We used the steps from the P/n protocol, except for a few cells with no P/n protocol, where we used a −10 mV test pulse before the axial current recording. We then correct for the loss in driving force due to imperfect clamping as in Traynelis (1998):

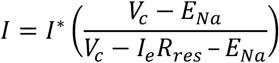

where *V_c_* is the command potential. In practice, this was a minor correction.

Figure 11B shows a recording from a retinal ganglion cell (black). After correction, the peak current is larger (red). We tested the effect of series resistance and the correction in a simple biophysical model of a RGC with an extended AIS, with the electrode modeled as a resistance (see *Biophysical Model*). Figure 11C shows that the peak recorded current decreases substantially when increasing *R_s_*, but this error is well corrected by the method described above. The spike threshold is only marginally affected by the series resistance (Fig. 11D; the exact value of the threshold depends on model parameters). We selected cells with residual series resistance smaller than 5 MΩ (*n* = 20). There was a correlation between measured peak axial current *I_p_* and *R_s_* (Fig. 11E, Pearson correlation r = 0.43, p = 0.06), but it remained small. We observed no correlation between spike threshold and *R_s_* (Fig. 11F, p = 0.25, Pearson test). Therefore, the impact of *R_s_* on our measurements should be moderate.

**Figure 11.**
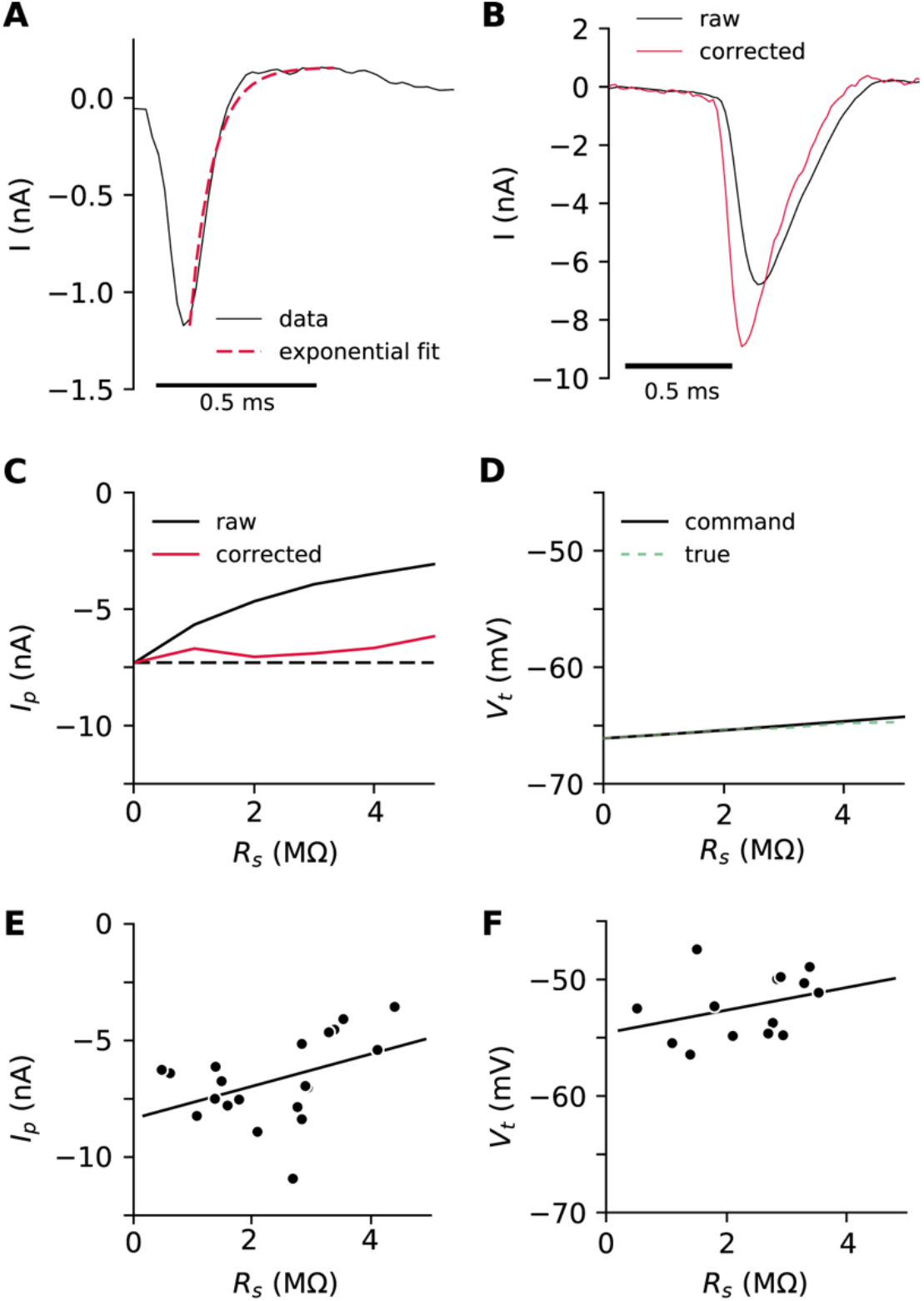
Correction of axial current recordings. A, Passive response to a +5 mV voltage step. An exponential function is fitted on the first 0.5 ms, starting from the peak, to estimate the decay time constant (dashed red). B, A recorded axial current (black) is corrected by defiltering (red), with residual Rs = 2.1 MΩ and τ = 70 μs. C, Peak current correction in a biophysical model with an extended AIS (5 to 30 μm from the soma), as a function of series resistance Rs. Dashed line: peak current in the ideal situation with no series resistance. D, Change in measured and actual voltage threshold as a function of Rs in a biophysical model. E, Corrected peak axial current at spike initiation vs. series resistance in RGCs (n = 20), showing a weak correlation (Pearson correlation r =0.43, p = 0.06). F, Corrected voltage threshold vs. Rs in RGCs (n = 14; 6 cells excluded because the reference potential drifted), with the regression line (p = 0.25, Pearson test).

#### Threshold

The voltage threshold *V_t_* is defined as the highest command potential where no axonal spike is elicited. The membrane potential at the soma differs slightly from the command potential by −*R*_*res*_*I*_*e*_. As the axial current at threshold is about 100 pA, this error is smaller than 0.5 mV. 6 cells for which the reference potential drifted by more than 3 mV during the recordings were discarded from voltage threshold analyses.

As the current at threshold is small, we used only the recordings with P/n protocol (n = 15) to ensure accurate leak subtraction. Three additional cells were excluded from the analysis of threshold current because the recordings were either too noisy or with unstable baseline current. Thus, n= 12 cells were used. The current traces below threshold were smoothed with a sliding window (half-window size is 50 points, 1 ms) before peak detection (Fig. 5A). The threshold current was then measured as the largest peak current for the data points between *V_t_*-1mV and *V_t_*.

#### Charge and current duration

The charge *Q* transferred to the soma at spike initiation is estimated as the integral of *I_e_* in the time window where the current is greater than 10% of its peak value (to avoid integrating noise). Current duration *t_50_* is the duration during which the current is greater than 50% of the peak value.

#### Adaptation

Relations between *V_t_* and *V_0_*(Fig. 7A) were fitted to the theoretical formula for threshold adaptation with sodium channel inactivation (Platkiewicz & Brette, 2011; Fontaine *et al.*, 2014):

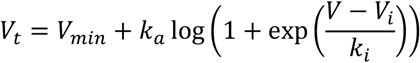

For the current adaptation analysis, cells were discarded if *R_s_* increased by more than 30% during the protocol. In a few cells, the threshold could not be clearly measured at −40 mV and therefore I_40_ is missing.

### Morphological data analysis

Axon tracing in 3D was performed automatically with Vaa3D (Peng *et al.*, 2010) (version 3.2 64-bit ran on MacOSX10.13.6) based on biocytin Z-stacks. The start and end position of the axon tracing was manually chosen by the researcher to include the entire AIS. Axon coordinates (*x*, *y*, *z*, radius) were stored in SWC files and analyzed with custom Python scripts. Coordinates were interpolated with a finer spacing corresponding to the pixel size (including in the *z* direction also). Interpolation was performed with B-splines using Scipy (Virtanen *et al.*, 2020) to evaluate the spline at each pixel comprised in the axon profile, and coordinates were rounded at the nearest pixel. The ankyrin-G images were then loaded as a 3D stack to get the fluorescence intensity at the interpolated coordinates along the axon profile. The intensity profile was smoothed with a sliding mean (half-width 15 pixels). The AIS start and end position were manually defined using the normalized intensity profile, the 3D stacks and the maximal intensity projection in Fiji (Schindelin *et al.*, 2012). Several cells for which the start or end position were considered too unclear to be determined accurately, were discarded from the analyses, so that morphological measurements were available for n = 14 cells.

### Statistics

The box plots display the distribution of data in the following way. The central bar shows the median. The lower and upper limit of the box shows the first and third quartiles (Q1 and Q3), respectively. The lower and upper whisker bars show Q1-1.5 IQR (interquartile range, the range between Q1 and Q3) and Q3 + 1.5 IQR, respectively. The data points outside the whiskers are outliers and indicated by diamonds marker.

### Theory

#### Axial current at spike initiation

The axial current at spike initiation has been derived previously based on resistive coupling theory (Hamada *et al.*, 2016):

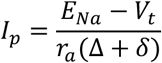

with

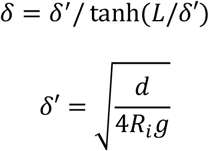

where Δ is AIS distance from the soma, *L* is AIS length, *d* is AIS diameter, *g* is Na^+^ conductance density, *R_i_* is intracellular resistivity and

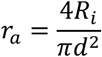

is axial resistance per unit length. The derivation makes the following assumptions: all Na^+^ channels are open; channel kinetics are neglected; capacitive and leak currents are considered negligible. These assumptions all tend to overestimate the axial current, but the approximation is generally good (see Fig. 4B). When *L* is much greater than δ′ (which was the case in our measurements), δ ≈ δ′ and the axial current is essentially insensitive to *L*. Intracellular resistivity has not been measured directly in the RGC axons. In dendrites of cortical pyramidal cells, it was estimated to be *R_i_* = 70-100 Ω.cm (Stuart & Spruston, 1998). Modeling studies in RGCs assume somewhat higher values, up to about 150 Ω.cm (Sheasby & Fohlmeister, 1999; Fohlmeister *et al.*, 2010), but these are based on model optimization. We chose *R_i_* = 100 Ω.cm (higher values would yield a higher estimate of Na^+^ channel conductance density).

The maximum current across all possible AIS geometries can be calculated by setting Δ =0 and *L* = ∞ (AIS of infinite length starting from the soma):

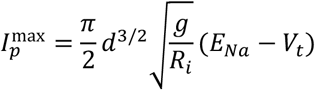

Therefore, the minimum conductance density necessary to produce an axial current *I* is:

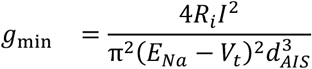

#### Axial current at threshold

In a model where the spatial extent of the AIS is neglected (all axonal Na^+^ channels clustered at a single point), the axial current at threshold is *k/R_a_*, where *k* is the Boltzmann activation slope of Na^+^ channels and *R_a_* is the axial resistance between soma and AIS (Brette, 2013). It is possible to calculate this current for an AIS of length *L* starting from the soma.

The axial current at threshold is:

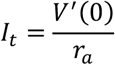

where *r_a_* is resistance per unit length. To obtain *V’(0)*, we solve the cable equation in a simple axon model where only the axial current and the Na^+^ current are considered, as in (Goethals & Brette, 2020). We consider a cylindrical axon of diameter *d*. The AIS has length *L* and starts from the somatic end. It has a uniform density of Nav channels. The total Nav conductance is

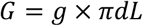

where *g* is the surface conductance density. We neglect leak and K^+^ currents, Nav channel inactivation, as well as all time-varying phenomena. The cable equation then becomes:

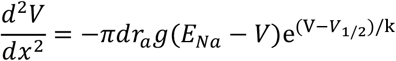

where *V*_1/0_ is the half-activation voltage of Nav channels. The boundary conditions are *V*(0) = *V_s_* (somatic potential) and *V’(L) = 0* (no axial current flowing towards the distal axon). In units of the AIS length *L*, this equation reads approximately:

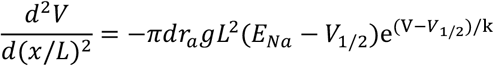

Here the driving force (*E_Na_* − *V*) has been approximated by (*E_Na_* − *V*_1/2_) as in (Brette, 2013). We now write the following change of variables:

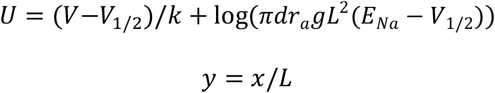

and we note *U*′ = *dU*/*dy*. That is, voltage is in units of *k* and space is in units of AIS length *L*. The rescaled cable equation is:

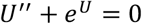

with the boundary conditions:

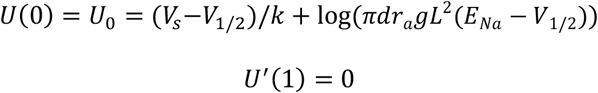

This equation is analytically solvable, with general solution

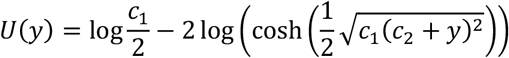

From *U*′ = 0, it follows that *c*_2_ = −1. We then obtain for the boundary condition at 0:

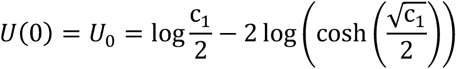

which defines c_1_ as an implicit function of *U*_0_. We look for a bifurcation, that is, a value of *U_0_* when the number of solutions changes. This is obtained by setting the derivative of the right hand-side to 0, which gives:

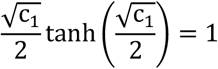

The solution can be calculated: √c_1_/2 ≈ 1.2, giving c_1_ ≈ 5.8. We have 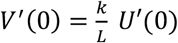 and

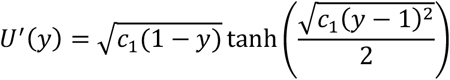

so *U*′(0) = 2 and we obtain:

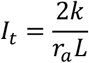

A simple extrapolated formula consistent with the formulae for both point AIS away from the soma and extended AIS starting from the soma is:

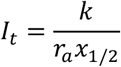

where *x*_1/2_ = Δ + *L*/2 is the middle position of the AIS, relative to the soma. In simulations, we find that this is a good approximation (Fig. 12).

**Figure 12.**
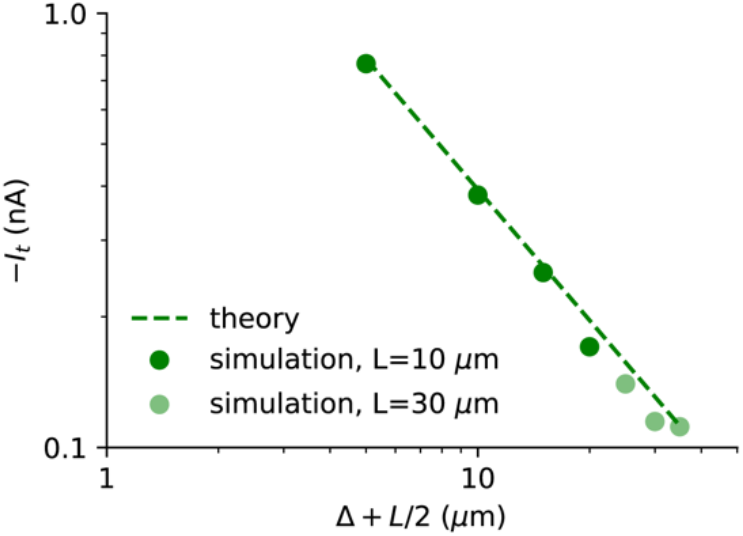
Theoretical estimation of axial current at threshold compared to simulations. The simple model (no inactivation, no Kv channels) is simulated with L = 10 μm (dark green dots) and L = 30 μm (light green dots), and the start position is varied from 0 to 20 μm. The theoretical curve is shown as a dashed line.

#### Axial current near threshold

We calculate the axial current just below threshold as a function of somatic voltage in a point AIS model. We consider a cylindrical axon of diameter *d* where all the Nav channels are located at a single location. The AIS contains a total Na^+^ conductance *G*. The axial current is

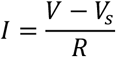

where *V* is the axonal voltage, *V_s_* is the somatic voltage, and *R* is the axial resistance between the soma and the AIS. It must equal the Na^+^ current:

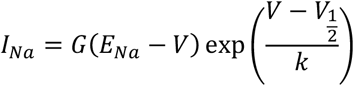

which is the exponential approximation near threshold. Near threshold, we have (*E_Na_* − *V*) ≈ (*E_Na_* − *V*_*s*_). We consider this driving force as a constant Δ*V*. We then absorb *V*_1/2_ into *G* and take *k* as the units of voltage. Thus, the equation reads:

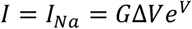

Using *V* = *V*_*s*_ + *RI*, we obtain:

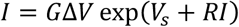

At the bifurcation (threshold), we have (differentiation with respect to *I*):

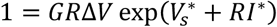

where 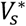 is the somatic voltage threshold and *I*\* is the axial current at threshold. We divide the two previous equations and obtain:

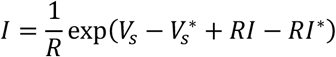

In a point AIS, the axial current at threshold is:

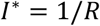

Therefore:

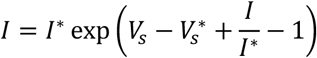

This can be rewritten as:

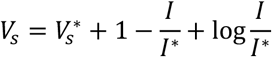

A Taylor expansion gives:

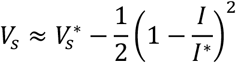

In original voltage units, we then obtain:

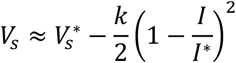

#### Relation between voltage threshold and axial current at spike initiation

Theoretically, voltage threshold varies as −*k* log *g*, where *g* is the available Na^+^ conductance (Platkiewicz & Brette, 2011; Brette, 2013; Goethals & Brette, 2020). The axial current at spike initiation also depends on *g*, and therefore voltage threshold and axial current co-vary when *g* is varied. The general relation is complicated, but a simple approximated relation can be obtained by considering the equation for the maximum current 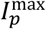. Since 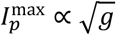, it follows that with this approximation the threshold varies as 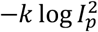, i.e., as −2*k* log *I*_*p*_.

### Simplified model

In Fig. 4A-B and Fig. 12, we used a simplified model with only non-inactivating Nav channels to check analytical expressions, similar to Brette (2013). A spherical soma of diameter 30 μm is attached to an axonal cylinder of diameter 1 μm and length 500 μm (soma diameter is in fact irrelevant as the soma is voltage-clamped). Specific membrane capacitance is *C_m_* = 0.9 μF/cm^2^; specific membrane resistance is *R_m_* = 15 000 Ω.cm2; leak reversal potential is *E_L_* = −75 mV; intracellular resistivity is *R_i_* = 100 Ω.cm. If not specified, Nav channels are placed from 5 μm to 35 μm on the axon. In Fig. 12, the length ranges from 10 μm to 30 μm and the start position from 0 μm to 20 μm. We used simple single gate activation dynamics with fixed time constant:

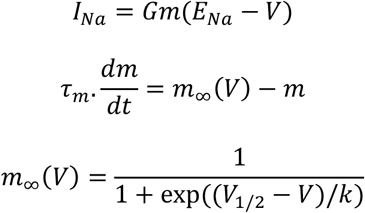

where *E_Na_* = 70 mV, *k* = 5 mV, *V_1/2_* = −35 mV and *τ_m_* = 53.6 μs (corresponding to 150 μs before temperature correction, see (Goethals & Brette, 2020)). For Fig. 4A and B, Na^+^ conductance density was *g* = 5000 S/m^2^. For Fig. 12, the total Na^+^ conductance was fixed (*G* = 350 nS) to keep the total number of Na^+^ channels fixed when AIS length is varied. This corresponds to conductance densities of about 11 100 and 3700 S/m^2^ for a 10 and 30 μm long AIS, respectively. The model is simulated in voltage-clamp and the threshold is measured with the bisection method. We used the Brian 2 simulator (Stimberg *et al.*, 2019) with 10 μs time step and 1 μm spatial resolution.

### Biophysical model

In Fig. 5B, 5D, 6E, 11C and 11D, we used a biophysical model of an AP with inactivating Nav channels and non-inactivating Kv channels, similar to (Goethals & Brette, 2020). The biophysical model has a simple geometry, consisting of a spherical soma (30 μm diameter), a long dendrite (diameter: 6 μm, length: 1000 μm) and a thin unmyelinated axon (diameter: 1 μm, length; 500 μm). The dendrite is irrelevant to most simulations because the soma is voltage-clamped, electrically isolating the dendrites from the axon. It only contributes an additional somatodendritic capacitance when an electrode model is added (Fig. 2). When not specified, the AIS extends from 5 μm to 35 μm from the soma. Specific membrane capacitance is *C_m_* = 0.9 μF/cm^2^; specific membrane resistance is *R_m_* = 15 000 Ω.cm^2^; leak reversal potential is *E_L_* = −75 mV; intracellular resistivity is *R_i_* = 100 Ω.cm.

In Fig. 5B and 5D, the AIS start position was 10 μm, close to the mean AIS start position in our cell population. The threshold was approached with 0.01 mV precision using the bisection method. In Fig. 6E, the AIS start position was varied from 0 to 20 μm and the AIS length was 30 μm. In these three panels, the Na^+^ conductance density was *g* = 3700 S/m^2^.

In Fig. 11C-D, we inserted an electrode model, which consists of a resistance *R_s_* (0 to 5 M Ω) between the amplifier and the soma, such that a current *(V_c_-V)/R_s_* is injected into the soma (where *V_c_* is the voltage command). The Na^+^ conductance density *g* was 7400 S/m^2^, to obtain peak axonal currents and thresholds comparable to measurements in RGCs.

**Table 1.**
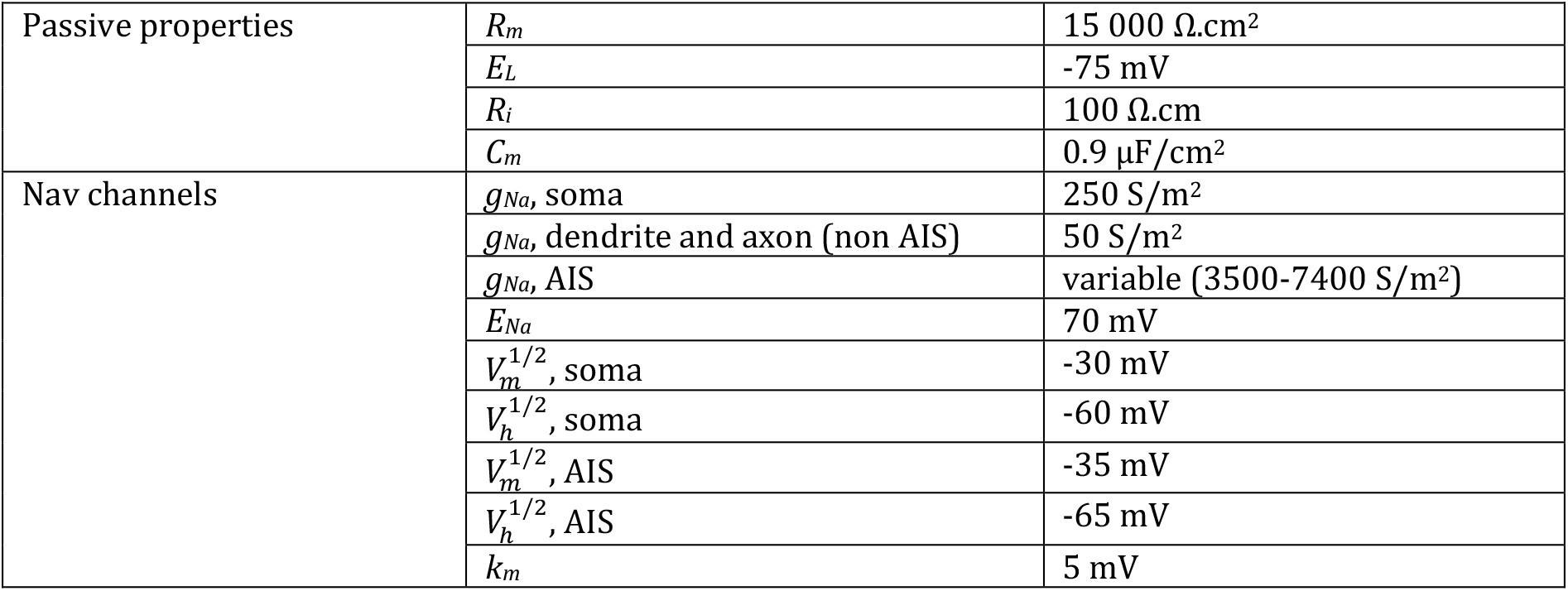
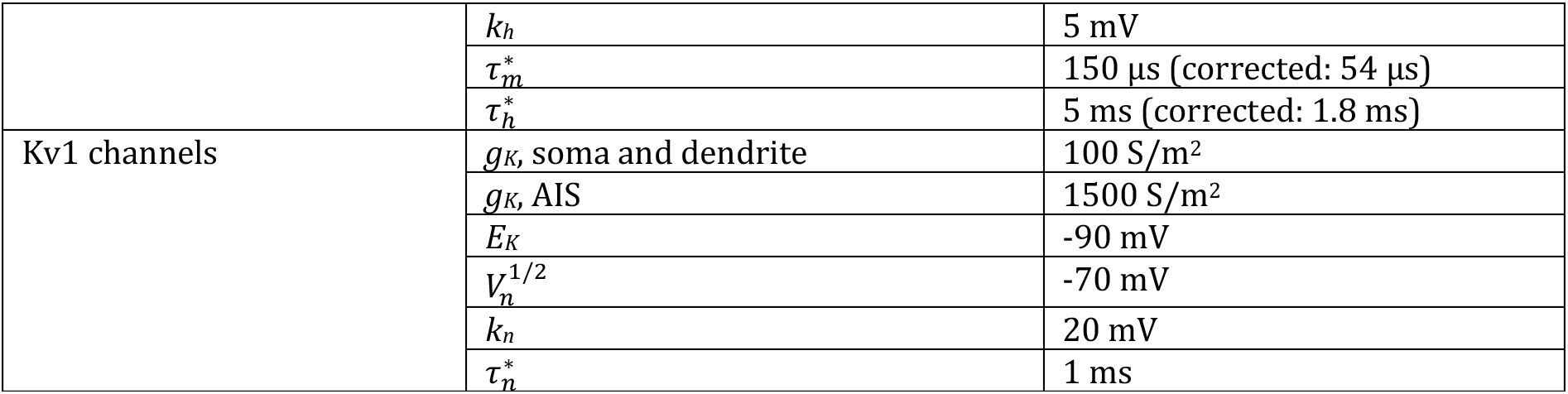
Parameters values of the biophysical model. Time constants corrected for temperature are indicated in brackets.

## Acknowledgments

We thank Marcel Stimberg for assistance with simulations, Elaine Orendorff for assistance with immunochemistry, Stéphane Fouquet for advice and assistance with confocal microscopy, Serge Picaud for access to the electrophysiology setup, and Boris Barbour and Christophe Leterrier for discussions.

